# Angiotensin blockade enhances motivational reward learning via enhancing striatal prediction error signaling and frontostriatal communication

**DOI:** 10.1101/2022.03.14.484364

**Authors:** Ting Xu, Xinqi Zhou, Jonathan W. Kanen, Lan Wang, Jialin Li, Zhiyi Chen, Ran Zhang, Guojuan Jiao, Feng Zhou, Weihua Zhao, Shuxia Yao, Benjamin Becker

## Abstract

Adaptive human learning utilizes reward prediction errors (RPEs) that scale the differences between expected and actual outcomes to optimize future choices. Depression has been linked with biased RPE signaling and an exaggerated impact of negative outcomes on learning which may promote amotivation and anhedonia. The present proof-of-concept study combined computational modelling and multivariate decoding with neuroimaging to determine the influence of the selective competitive angiotensin II type 1 receptor antagonist losartan on learning from positive or negative outcomes and the underlying neural mechanisms in healthy humans. In a double-blind, between-subjects, placebo-controlled pharmaco-fMRI experiment, 61 healthy male participants (losartan, n=30; placebo, n=31) underwent a probabilistic selection reinforcement learning task incorporating a learning and transfer phase. Losartan improved choice accuracy for the hardest stimulus pair via increasing expected value sensitivity towards the rewarding stimulus relative to the placebo group during learning. Computational modelling revealed that losartan reduced the learning rate for negative outcomes and increased exploitatory choice behaviors while preserving learning for positive outcomes. These behavioral patterns were paralleled on the neural level by increased RPE signaling in orbitofrontal-striatal regions and enhanced positive outcome representations in the ventral striatum (VS) following losartan. In the transfer phase, losartan accelerated response times and enhanced VS functional connectivity with left dorsolateral prefrontal cortex when approaching maximum rewards. These findings elucidate the potential of losartan to reduce the impact of negative outcomes during learning and subsequently facilitate motivational approach towards maximum rewards in the transfer of learning. This may indicate a promising therapeutic mechanism to normalize distorted reward learning and fronto-striatal functioning in depression.

## INTRODUCTION

Human learning is driven by reward prediction errors (RPEs) that signal the discrepancy between expected and actual outcomes. Computational approaches have closely linked RPEs to dopaminergic signaling in the midbrain-striatum circuitry and to motivation and reward seeking [1, 2]. Deficits in these domains, in particular amotivation and anhedonia, represent key symptoms of unipolar depression and dysregulated RPE signaling has been proposed as a potential underlying neurocomputational candidate mechanism [3]. Specifically, within a computational reinforcement learning (RL) framework, depressed individuals showed enhanced sensitivity to negative information while they concomitantly discounted positive feedback leading to reduced learning from positive events [4, 5]. On the neural level this learning bias was often accompanied by blunted RPE signaling in the ventral striatum and reduced fronto-striatal connectivity during reward feedback [6, 7]. These neural dysregulations have been associated with depressive symptom load, specifically anhedonia and persistent negative mood [8], and could predict anti-depressive treatment responses [9]. As such, distorted learning from negative and positive outcomes may play a key role in the pathophysiology of depression and may represent a promising target for novel antidepressive treatments.

Accumulating evidence suggests that the renin-angiotensin system (RAS) plays a key role in learning. Preclinical work in rodents and humans has utilized the selective competitive angiotensin II type 1 receptor (AT1R) antagonist losartan (LT) - an approved treatment for hypertension with an excellent safety record [10, 11] - to modulate learning from negative or positive events [12, 13]. Recent human studies have demonstrated that a single dose of LT selectively suppressed memory encoding of threatening materials [14] and accelerated threat extinction learning [15, 16]. Moreover, LT specifically affected probabilistic learning from negative outcomes by reducing the degree to which participants learned from loss feedback, while leaving learning from positive outcomes unaffected [12]. An initial neuroimaging study moreover reported modulatory effects of LT on mesocorticolimbic functional connectivity during social reward and punishment processing [17].

Given the pivotal role of dopamine (DA) in RPE signaling and modulation of the mesocorticolimbic circuits [18], these results may indicate a downstream effect of LT on DA signaling and in turn on learning from positive and negative outcomes. Support for a potential DA-mediated mechanism of action is provided by studies suggesting an important role of the AT1R in regulating central dopaminergic neurotransmission [19], high co-expression of AT1R and DA receptors [20] and evidence for functionally interactions between AT1R and DA receptors in the striatum [21].

Against this background, the present proof-of-concept study combined computational modelling and functional MRI, with a preregistered between-subjects randomized double-blind placebo-controlled pharmaco-fMRI design in n = 61 [Male] healthy participants to determine modulatory effects of LT-induced AT1R blockade on RL model parameters and the underlying neural mechanisms. We utilized a validated probabilistic selection RL paradigm with two stages: a learning phase in which participants learned to make better choices for fixed pairs of stimuli according to reward or loss feedback, and a subsequent transfer phase in which participants applied the learned optimal choices to novel combinations of stimuli without feedback. Behavioral responses during learning were fit using a computational RL model to describe the dynamic learning process. Effects of LT on learning were examined by means of comparing RL model parameters and neural activity related to model-derived estimates of RPE. Effects of LT on learning transfer were examined by comparing choices and functional connectivity in cortico-striatal pathways when participants approached the best or avoided the worst stimulus. Based on previous literature [14, 16, 17], we predicted that LT would: 1) reduce learning from negative outcomes, increase the RPE-associated signaling in the ventral striatum (VS) and its neural expression for positive outcomes during the learning phase and 2) increase selection of the best stimulus in the context of increased fronto-striatal coupling during learning transfer.

## METHODS

### Participants

Seventy right-handed healthy male Chinese participants were screened according to evaluated enrollment criteria (see **Supplemental Methods**, sample size based on previous studies [15-17]). The study focused on male individuals to control for sex differences in response to RAS blockade [22] and menstrual cycle-dependent variations in reward processing [23]. Nine participants exhibited excessive head movement (LT, n=1) or poor learning (LT, n=4; Placebo, n=4, **Fig. 1**) and thus failed to reach the choice accuracy criteria (i.e., choosing 65% A in AB, 55% C in CD, 50% E in EF, in line with Frank et al., 2007), leading to a final sample of n = 61 (mean±SD, age=20.89±2.32 years).

**Fig. 1.**
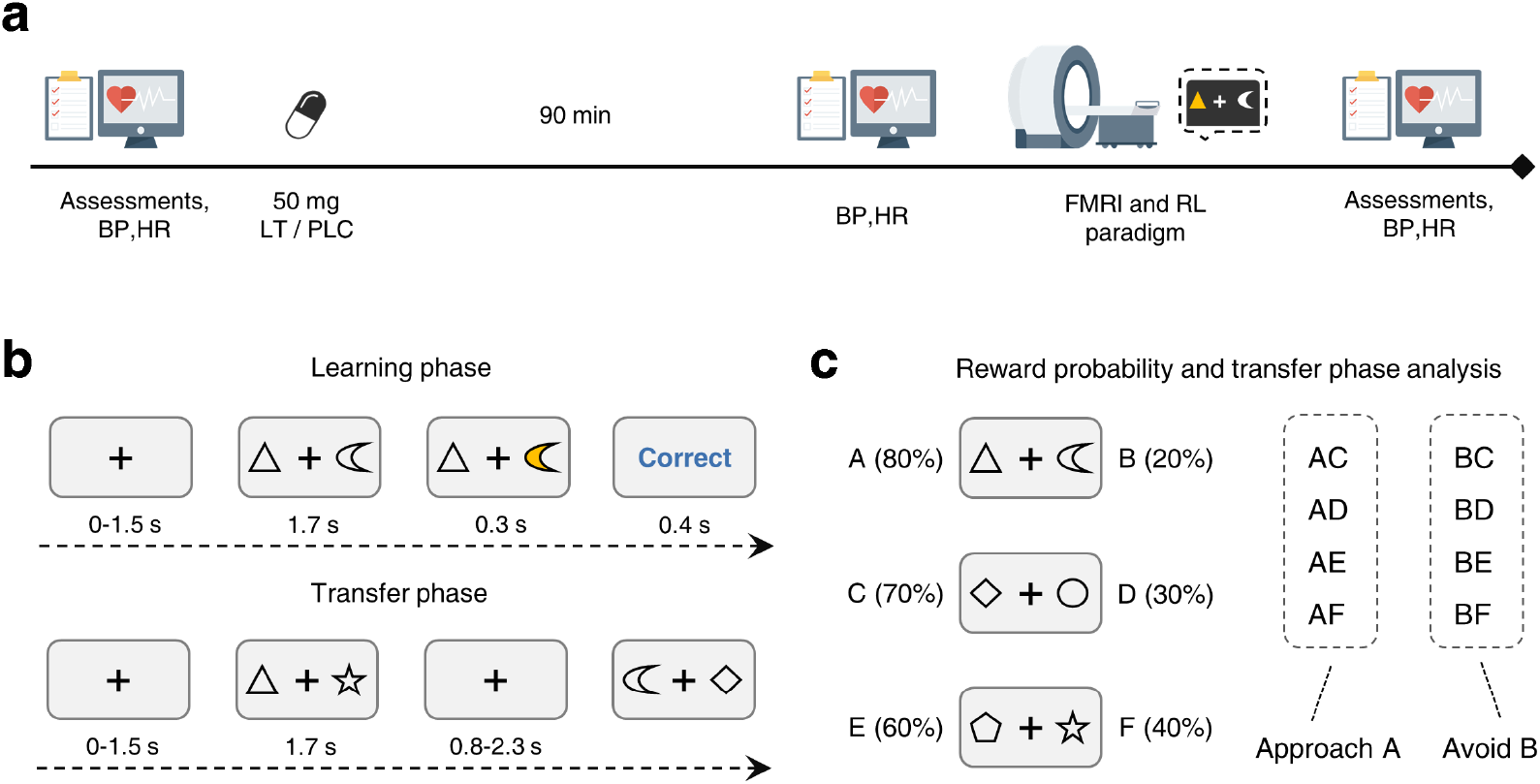
Experimental timeline and paradigm. **(a)** Experimental timeline. (**b)**. The reinforcement learning paradigm consisted of two subsequent phases. During the learning phase participants were presented with one of three different pairs of six stimuli (denoted as AB, CD and EF) on each trial in a randomized order. Participants were instructed to learn to choose the best option within each stimulus pair based on the feedback presented (i.e., ‘correct’ or ‘wrong’ presented as text, which indicated that 0.5 RMB or nothing were added to the total payment). To avoid choice preference or reward associations with one particular stimulus, stimulus pairs were presented in a counterbalanced order across subjects. During the transfer phase, participants were presented with all permutations of combinations with A and B - corresponding to the stimuli with the highest or lowest reward probability, respectively - and were instructed to choose the better option according to their previous learning experience. **(c)** The probabilities of acquiring reward for pairs AB, CD and EF were 80:20, 70:30 and 60:40, respectively, during the learning phase. The easiest condition was therefore the AB pair while the EF pair was the hardest one to learn because of the relatively equivalent reward probabilities between the two stimuli. A performance criterion (i.e., choosing 65% A in AB, 55% C in CD, 50% E in EF, a similar approach to that used by Frank et al., 2007) was initially used to ensure successful learning in the subjects. N = 4 subjects in each treatment group did not fulfill that criterion and were excluded from the further analysis. The analysis in the transfer phase was conducted on trials in which A was correctly chosen or B was avoided when being paired with another stimulus. Abbreviations: BP, blood pressure; HR, heart rate; LT, losartan; PLC, placebo; FMRI, functional magnetic resonance imaging; RL, reinforcement learning. License: Image in 1a were designed by DinosoftLabs and obtained from Flaticon.com under the free use license.

All participants provided written informed consent, protocols were pre-registered on Clinical Trials.gov (https://clinicaltrials.gov/ct2/show/NCT04604938) and approved by the ethics committee at the University of Electronic Science and Technology of China (Approval 355).

Using a double-blind randomized, placebo-controlled, between-subjects pharmacological fMRI design, participants were administered either a single oral dose of LT (50mg) or placebo (PLC) packed in identical capsules. Capsules were dispensed by an independent researcher based on a computer-generated randomization sequence to implement double blinding. Investigators involved in data acquisition and analyses were blinded for group allocation. Prior studies suggest that single-dose LT effects on cardiovascular activity in healthy subjects were observed after 3 hours [24]. Effects of the blood-brain barrier permeable LT on brain activity and cognitive domains were observed in the timeframe ranging from 1.5-2.5h after single-dose administration (which overlaps with peak plasma levels after 90 minutes and an elimination half-life between 1.5-2.5h [25-27]). Consistent with this, LT treatment was administered 90min before fMRI acquisition. Participants first performed a reinforcement learning task (duration 30min) followed by an emotional memory task (reported in Xu et al., 2021 [14]). To control for nonspecific effects of LT, assessments of mood, attention and memory were incorporated at baseline and after the experiment, while cardiovascular activity (i.e., blood pressure, heart rate) was measured at baseline, after drug administration and after the experiment (**Fig. 1a** and **Supplemental Methods**). To ensure double blinding participants were asked to guess treatment after the experiment (treatment guess c^2^=0.40, p=0.53; confirming successful double-blinding).

## Experimental design

### Probabilistic selection reinforcement learning paradigm

A validated probabilistic selection reinforcement learning paradigm was employed [28-30]. This paradigm consisted of two stages: an initial reinforcement learning phase and a subsequent transfer phase. During the learning phase, participants were presented with one of three different pairs of six shape stimuli (denoted as AB, CD and EF, **Fig. 1c**) on each trial in a randomized order. Participants were instructed to learn to choose the better option of each stimulus pair based on feedback (**Fig .1b**). Learning difficulty varied for the stimulus pairs in terms of reward contingency (80:20, 70:30 or 60:40, for AB, CD or EF, respectively). A total of 240 trials – dispersed across two fMRI runs with 120 trials each (40 trials per stimulus pair, trial mean duration 4s) – were presented during the learning phase. Each trial began with a fixation cross presented for a jittered interval of 0, 500, 1000, or 1500ms (**Fig. 1b**) followed by the presentation of two shapes displayed to the left and right of the fixation cross (side was counterbalanced). Stimuli were presented until participants made a response or 1700ms elapsed. The choice was visually confirmed by highlighting the chosen shape in yellow for 300ms, followed by 400ms feedback presentation (‘correct’ or ‘wrong’). Then, the fixation cross was displayed again until the whole trial duration was reached. In addition, 12 null trials without stimulus presentation of the same duration were randomly interspersed in each fMRI run to improve the model fitting of the rapid event-related fMRI design.

For the transfer phase, the six shape stimuli were recombined to constitute fifteen stimulus pairs. Each stimulus pair permutation was presented 8 times (side was counterbalanced) leading to 120 trials in the transfer phase, also with 12 null trials interspersed in a random order. Duration of each trial was 1700ms and no feedback was provided (**Fig. 1b**). Participants were told to choose the better option in each stimulus pair according to what they had learned in the learning phase.

### Computational modeling of learning behavior

We explored the learning rate in terms of choice behavior by using a Q-learning algorithm [31]. The Q-learning algorithm has been widely employed to model learning behavior and serves to model the change in choice behavior based on trial-by-trial updates of the expected value of choice options [30, 32, 33]. The corresponding model contains three free parameters: learning rate for positive (αGain) and negative (αLoss) RPEs and estimation of explore-exploit tendency (β). For details of modeling procedures, model estimation and comparison see **Supplemental Methods and Fig. S1**.

### Statistical analyses on the behavioral level

All analyses were performed in R (R development core and team, 2017). For the learning phase, we employed a multilevel Bayesian linear model to analyze trial-by-trial choice behavior using the Bayesian regression model in Stan (brms) R package [34]. Main effects of treatment, stimulus pair and fMRI run, as well as the interaction of stimulus pair and treatment on choice accuracy (proportion of choosing better stimulus in one stimulus pair, e.g., choosing A in AB pair) were considered as credibly different when more than 95% of the posterior distribution was above/below zero. Treatment effects on computational modeling indices of choice behavior (learning rate, explore-exploit tendency) were examined by using permutation-based two sample t tests via functions in the user contributed packages permuco (version 1.1.0) for the R environment.

In the transfer phase, we performed similar analyses for trials including stimuli with the highest and lowest reward probability (A, 80% or B, 20%) (**Fig. 1c**) to examine LT effects on choosing the best and avoiding the worst option. An exploratory model additionally examined effects of LT on choice times with choice (choose A, avoid choosing B) and treatment (LT, PLC) as fixed factors and subject as random factor. Main effects of treatment, choice behavior and their interaction were considered significant using the same 95% posterior distribution criterion (details see **Supplemental Methods**). We additionally explored the effects of treatment and rewarding probability (value) difference (e.g., AB-60, AD-50, AF-40, CF-30, AE-20, AC-10) on choice accuracy and reaction time during learning transfer (details see **Supplemental Methods**) to control for a potential confounding impact of probability difference levels on treatment effects. We did not find significant main effects of treatment or an interaction effect between treatment and rewarding probability difference (**Fig. S2, Table S1-S2**), suggesting rather specific treatment effects on easily-learned stimuli (i.e., A, B) during learning transfer.

### MRI acquisition, preprocessing and first level analysis

MRI data were acquired on a 3.0-T GE Discovery MR system (General Electric Medical System, Milwaukee, WI, USA) and preprocessed using standard procedures in SPM 12 (Statistical Parametric Mapping; http://www.fil.ion.ucl.ac.uk/spm/; Wellcome Trust Centre for Neuroimaging) (see **Supplemental Methods**).Separate general linear models (GLM) were designed for the learning and transfer phase.

For the learning phase, we established a GLM model that incorporated separate outcome onsets for positive and negative feedback, each modulated by the corresponding RPE estimated from the computational model. The highlight-period and six head motion parameters were all included as covariates of no interest. Given that the choice accuracy in both treatment groups rapidly approached a ceiling effect (such that irrespective of treatment participants rapidly showed optimal choice accuracy, i.e., >80%, across stimulus pairs with different rewarding probability in the second run, **Fig. S3**), the first and second fMRI run were modeled separately, and analyses focused on the first run to increase the sensitivity for learning-associated treatment effects.

During the transfer phase, approach A and avoid B were modeled as separate conditions, and the six head motion parameters were included as nuisance regressors.

### Examining neural effects of LT on RPE signaling during early learning

To examine the effects of LT on RPE signaling during early learning, the corresponding first level contrasts (i.e., Positive+Negative RPE) were subjected to voxel-wise two sample t tests. Whole brain analyses thresholded at cluster level family-wise error (FWE) corrected p<0.05 were employed (initial cluster threshold, p<0.001 uncorrected; see recommendations in Slotnick, 2017 [35]).

### Effects of LT on feedback-sensitive neural expressions in the VS during early learning

Given the higher sensitivity of multivariate neurofunctional representations for a given mental process including reward and RPEs in the VS [36, 37], multi-voxel pattern analysis (MVPA) was employed (see **Supplemental Methods**). We initially developed a decoder on the whole brain neural pattern that differentiated positive versus negative outcomes during early learning and tested it in an independent sample to validate brain systems strongly involved in differentiating reward versus loss. Next, treatment effects on the corresponding expression in the VS were examined (details see **Supplemental Methods**). The VS region of interest included the ventral caudate and nucleus accumbens, defined from the brainnetome atlas [38], which was functionally validated in our previous work [39, 40].

### Examining neural effects of LT on optimal choice behavior during learning transfer

Effects of LT on the transfer of optimal choice behavior were examined by means of separate voxel-wise two sample t-tests for choosing A (choosing the best option) or avoiding B (avoiding the worst option), respectively. We employed the whole brain analyses thresholded at cluster level family-wise error (FWE) corrected p<0.05 (initial cluster threshold, p<0.001 uncorrected).

### Functional connectivity analysis during learning transfer

Given that animal and human studies indicate that reinforcement learning is critically mediated by the functional communication between the VS and frontal regions [41, 42], we explored whether LT treatment could affect frontal-VS functional connectivity when subjects approach maximum (approach A) or avoid worst rewards (avoid B) during transfer phase. Treatment effects on frontal-VS functional networks were determined by performing two sample t tests on choosing A or avoid choosing B events. Within the brainnetome atlas-defined prefrontal cortex, results were thresholded at p<0.05 FWE corrected at peak level with small volume correction (SVC). In addition, we also explored the role of the VS in perceiving rewarding probability (value) difference by exploratory two sample t test with treatment as independent variable on VS activation for each value difference level.

## RESULTS

### Demographics and potential confounders

The LT (n=30) and PLC (n=31) groups were comparable with respect to sociodemographics and mood and cardiovascular indices arguing against nonspecific treatment effects (**Table 1**; all ps>0.10).

**Table 1.**
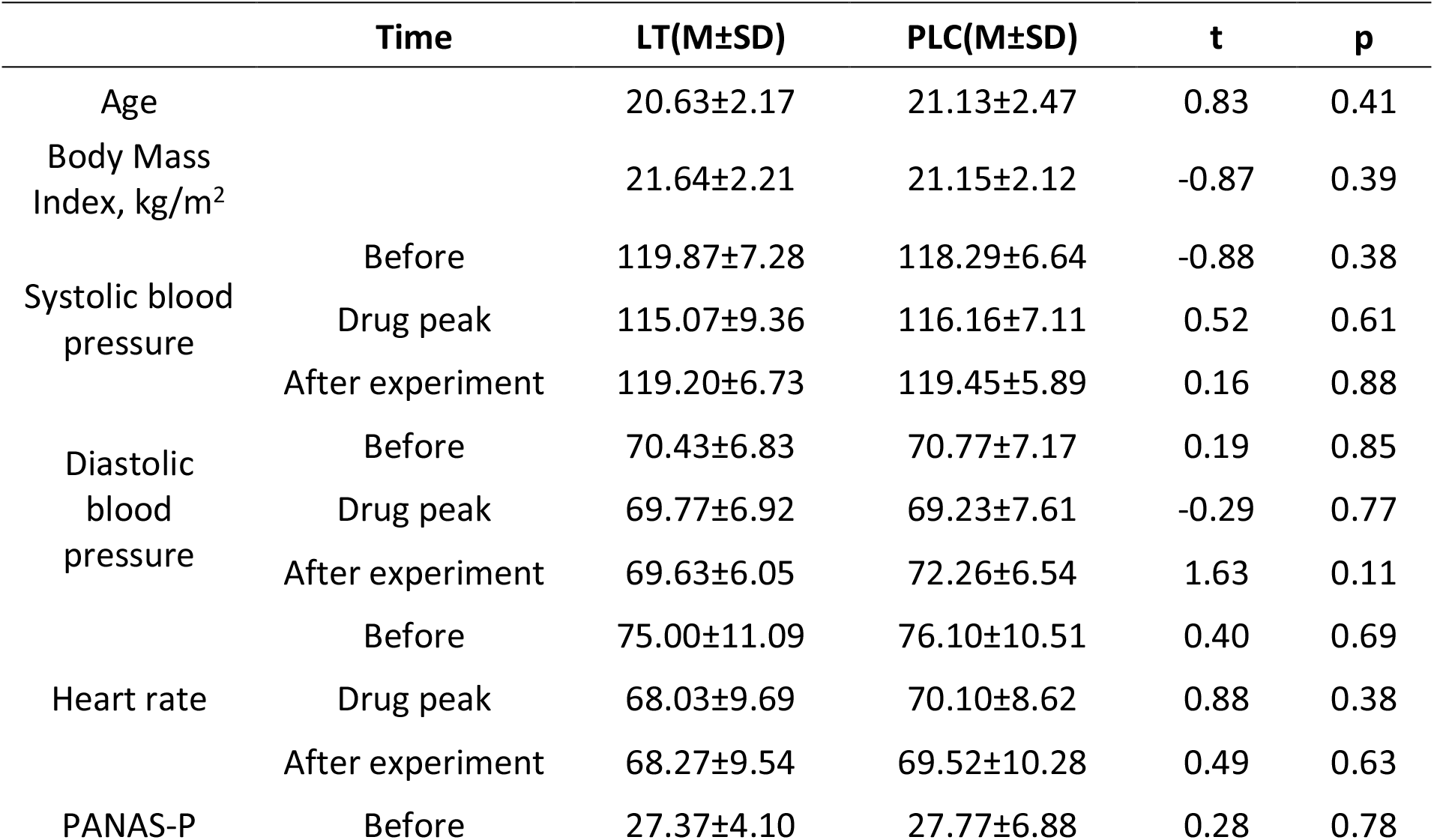
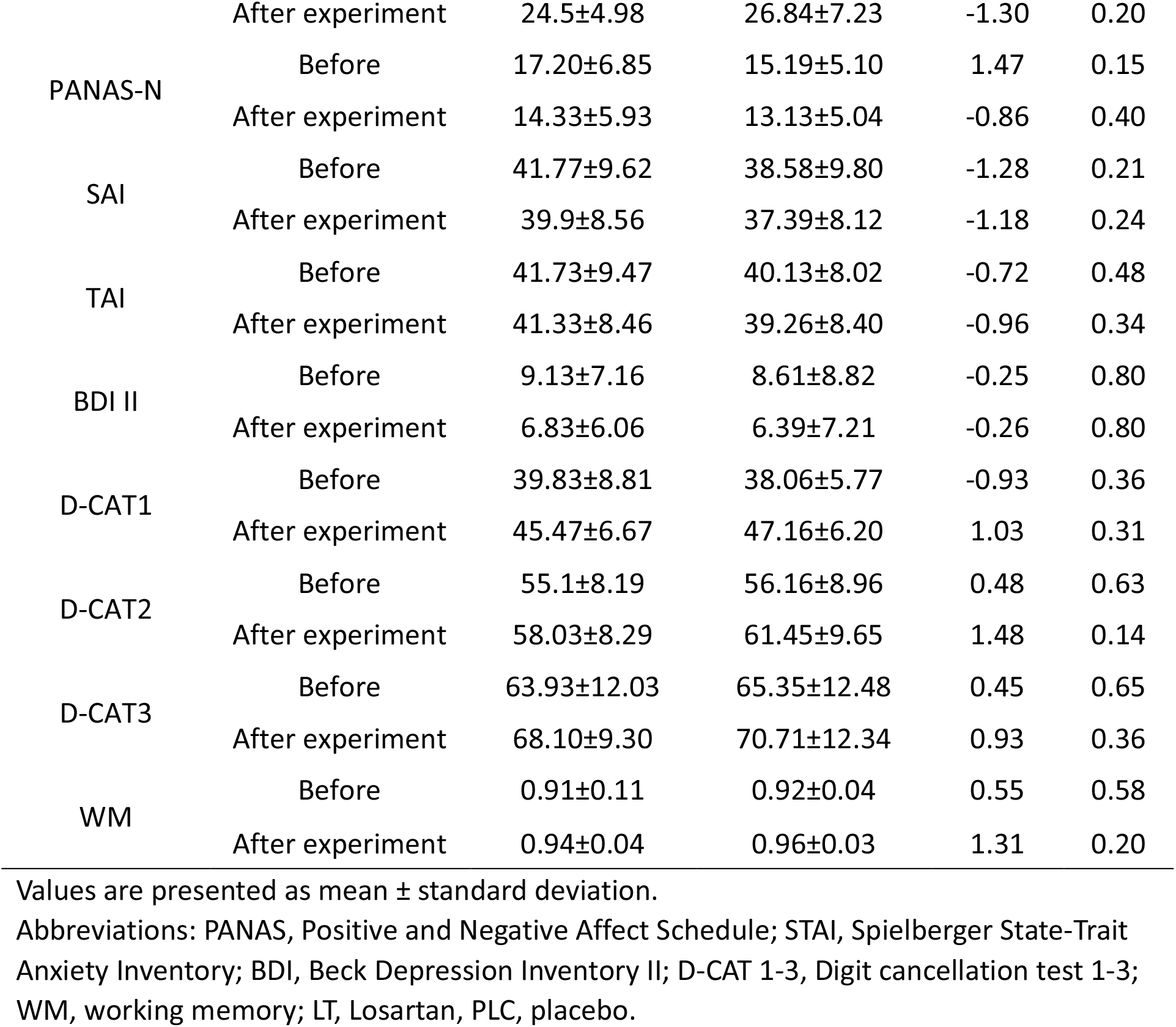
Demographics and potential confounders in two groups.

|  | Time | LT(M±SD) | PLC(M±SD) | t | p |
| --- | --- | --- | --- | --- | --- |
| Age |  | 20.63±2.17 | 21.13±2.47 | 0.83 | 0.41 |
| Body Mass Index, kg/m<sup>2</sup> |  | 21.64±2.21 | 21.15±2.12 | -0.87 | 0.39 |
| Systolic blood pressure | Before | 119.87±7.28 | 118.29±6.64 | -0.88 | 0.38 |
|  | Drug peak | 115.07±9.36 | 116.16±7.11 | 0.52 | 0.61 |
|  | After experiment | 119.20±6.73 | 119.45±5.89 | 0.16 | 0.88 |
| Diastolic blood pressure | Before | 70.43±6.83 | 70.77±7.17 | 0.19 | 0.85 |
|  | Drug peak | 69.77±6.92 | 69.23±7.61 | -0.29 | 0.77 |
|  | After experiment | 69.63±6.05 | 72.26±6.54 | 1.63 | 0.11 |
| Heart rate | Before | 75.00±11.09 | 76.10±10.51 | 0.40 | 0.69 |
|  | Drug peak | 68.03±9.69 | 70.10±8.62 | 0.88 | 0.38 |
|  | After experiment | 68.27±9.54 | 69.52±10.28 | 0.49 | 0.63 |
| PANAS-P | Before | 27.37±4.10 | 27.77±6.88 | 0.28 | 0.78 |
|  | After experiment | 24.5±4.98 | 26.84±7.23 | -1.30 | 0.20 |
| PANAS-N | Before | 17.20±6.85 | 15.19±5.10 | 1.47 | 0.15 |
|  | After experiment | 14.33±5.93 | 13.13±5.04 | -0.86 | 0.40 |
| SAI | Before | 41.77±9.62 | 38.58±9.80 | -1.28 | 0.21 |
|  | After experiment | 39.9±8.56 | 37.39±8.12 | -1.18 | 0.24 |
| TAI | Before | 41.73±9.47 | 40.13±8.02 | -0.72 | 0.48 |
|  | After experiment | 41.33±8.46 | 39.26±8.40 | -0.96 | 0.34 |
| BDI II | Before | 9.13±7.16 | 8.61±8.82 | -0.25 | 0.80 |
|  | After experiment | 6.83±6.06 | 6.39±7.21 | -0.26 | 0.80 |
| D-CAT1 | Before | 39.83±8.81 | 38.06±5.77 | -0.93 | 0.36 |
|  | After experiment | 45.47±6.67 | 47.16±6.20 | 1.03 | 0.31 |
| D-CAT2 | Before | 55.1±8.19 | 56.16±8.96 | 0.48 | 0.63 |
|  | After experiment | 58.03±8.29 | 61.45±9.65 | 1.48 | 0.14 |
| D-CAT3 | Before | 63.93±12.03 | 65.35±12.48 | 0.45 | 0.65 |
|  | After experiment | 68.10±9.30 | 70.71±12.34 | 0.93 | 0.36 |
| WM | Before | 0.91±0.11 | 0.92±0.04 | 0.55 | 0.58 |
|  | After experiment | 0.94±0.04 | 0.96±0.03 | 1.31 | 0.20 |

Values are presented as mean ± standard deviation.
Abbreviations: PANAS, Positive and Negative Affect Schedule; STAI, Spielberger State-Trait Anxiety Inventory; BDI, Beck Depression Inventory II; D-CAT 1-3, Digit cancellation test 1-3; WM, working memory; LT, Losartan, PLC, placebo.

### LT increases choice accuracy for the hardest stimulus pair via increasing value sensitivity during early learning

The choice accuracy indicates the proportion of trials on which subjects chose the option with higher probability for reward in a stimulus pair (e.g., choose A in AB). Here we observed a significant main effect of stimulus pair (β=0.19, 95% highest-density interval (HDI), [0.16, 0.22], **Fig. S4**) such that participants exhibited the highest choice accuracy for the easy stimulus pair. The main effect of treatment did not reach significance (β=0.02, 95% HDI, [-0.02, 0.06], **Fig. S4**), but the main effect of fMRI run (β=0.12, 95% HDI, [0.11, 0.13], **Fig. S4**) was significant. Further inspection revealed that participants in both treatment groups showed an increased learning trend with high (all>85.3%) choice accuracy across stimulus pairs with different rewarding probability (**Fig. S3**). This may reflect that factors not related to probabilistic learning per se such as an understanding of the task structure or the reward probabilities may have led to an improved learning performance in the second run irrespective of treatment. The ceiling effect will lead to biased estimates and could critically reduce the sensitivity of detecting treatment effects on the trial-wise feedback-dependent learning process under examination (see also [43] for treatment effects on between run learning performance). This pattern was confirmed by a robust treatment × stimulus pair interaction effect in the first (β=-0.05, 95% HDI, [-0.07,-0.03], **Fig. 2a**) but not the second fMRI run (β=0.01, 95% HDI, [-0.01, 0.03], **Fig. 2b**), with further analyses indicating that compared to PLC – LT increased choice accuracy for the most difficult stimulus pair (EF, β=0.09, 95% HDI, [0.01, 0.18], **Fig. 2c**), especially during the early learning period for that pair (trial 1-10, t_(59)_=3.55, p=0.01, trial 11-20, t_(59)_=7.29, p<0.001, **Fig. 2d**), but not the easier pairs (AB, β=-0.01, 95% HDI, [-0.07,0.05]; CD, β=0.03, 95% HDI, [-0.05, 0.10], **Fig. 2c**) during the first run. Our exploratory mediation analyses further indicated that Q value sensitivity only played a mediating role in the effects of LT on choice accuracy for the hardest stimulus pair (details see **Supplemental Methods and Fig. S5**). To increase the sensitivity to determine learning-related treatment effects, all subsequent behavioral and neural analysis consequently focused on the early learning phase (run 1).

**Fig. 2.**
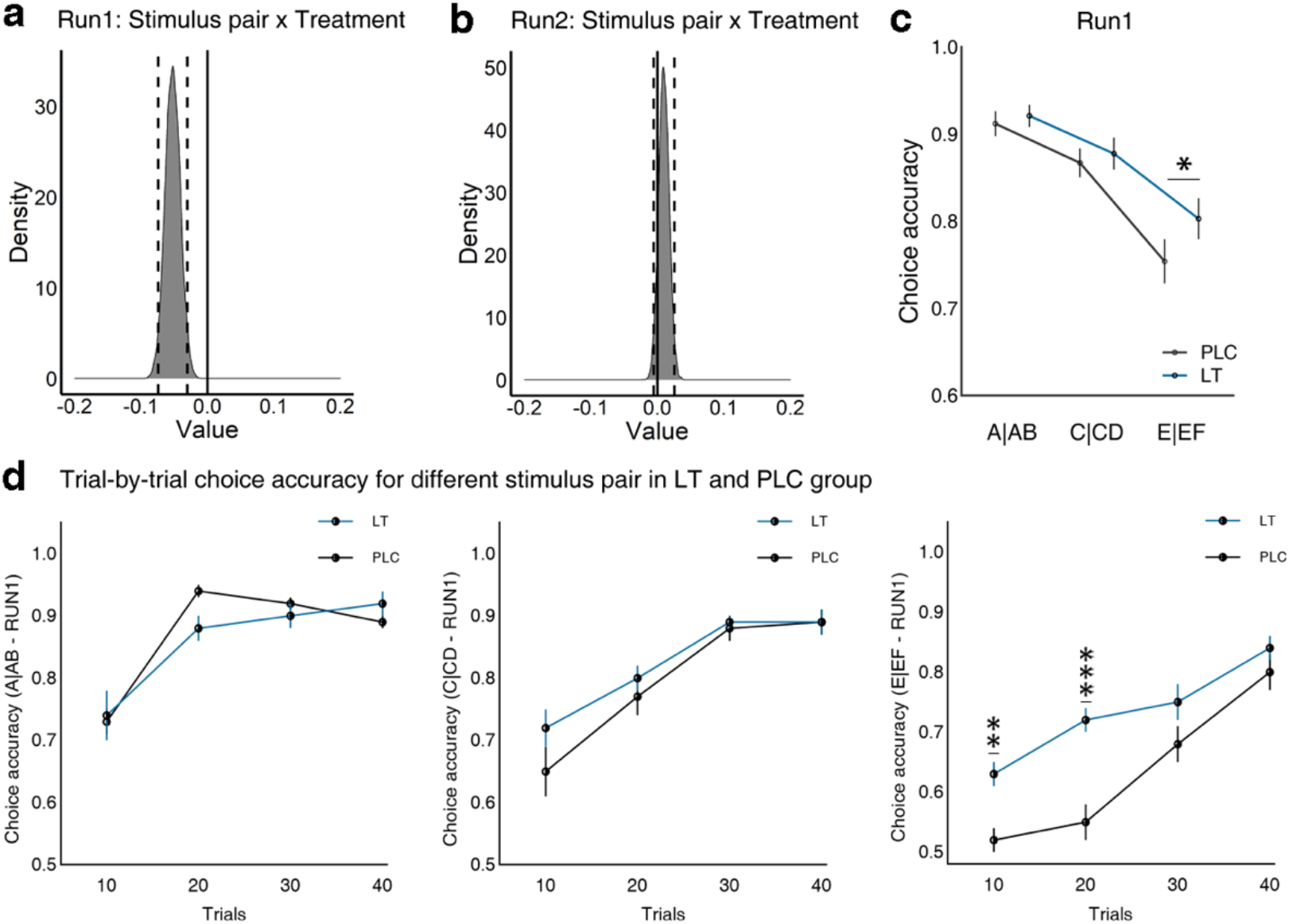
Behavioral effects of Losartan on choice accuracy and choice time. **(a-b)** A significant interaction between treatment and stimulus pair was observed in the first but not the second fMRI run of the learning phase. **(c)** During the first fMRI run of the learning phase, losartan-treated participants increased the choice accuracy for the hardest stimulus pair (EF) relative to the placebo group. **(d)** Specifically, in the first fMRI run, losartan group learned faster relative to placebo group during the initial 20 trials for the hardest stimulus pair. PLC, placebo; LT, losartan; *p<0.05, **p<0.01, ***p<0.001.

### LT reduces learning rate for negative outcomes during early learning

In line with our hypothesis, LT significantly reduced learning rate from negative outcomes (t_(59)_=-2.40, p=0.02, d=-0.61, **Fig. 3b**) but did not affect learning from positive outcomes (t_(59)_=-1.84,p=0.07,d=-0.47, **Fig. 3a**). Moreover, LT enhanced the explore-exploit parameter (t_(59)_=3.83, p<0.01, d=0.98, **Fig. 3c**), reflecting increased exploitatory choice behavior in terms of more consistent selection of options with higher expected reward values.

**Fig. 3.**
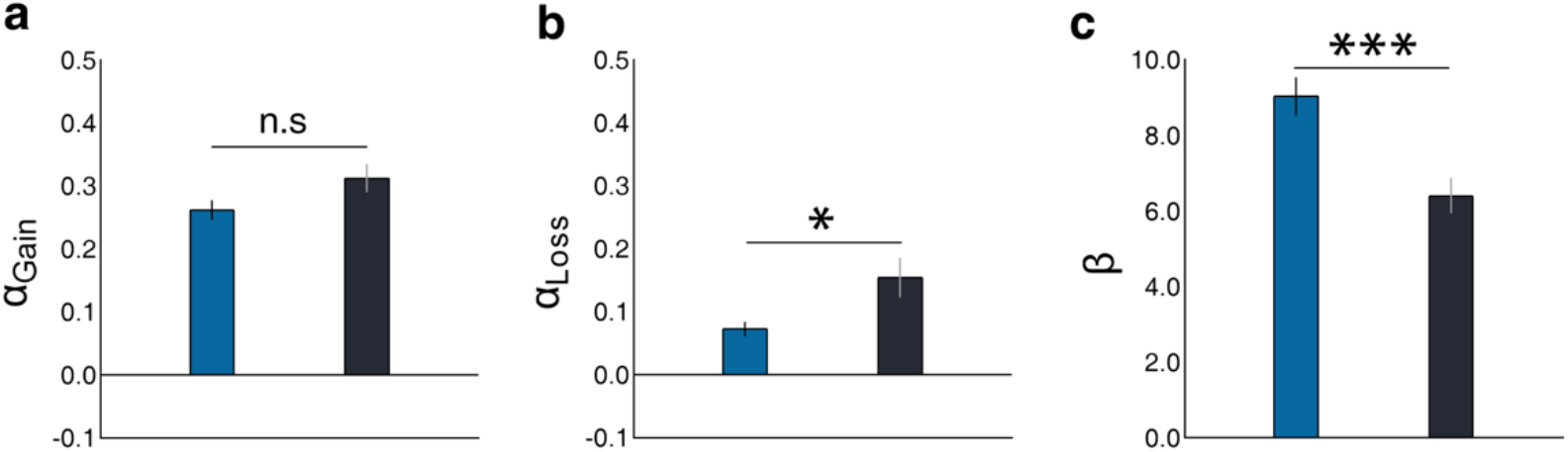
Losartan effects on computational model parameters. **(a)** Losartan and placebo groups showed equivalent learning rate for positive outcome. **(b)** Compared to the placebo group, losartan-treated individuals exhibited a reduced learning rate for negative outcomes. **(c)** Moreover, losartan enhanced exploitatory decisions relative to placebo group. The error bars denote standard error of the mean. n.s, non-significant, p<0.05, *** p<0.01.

### LT increases RPE signaling during early learning

We initially examined brain regions that scaled positive and negative RPEs independent of treatment. A corresponding one sample t-test confirmed previous studies suggesting that activity in striatal and frontal regions linearly increased with the strength of the RPEs (**Fig. S6, Table S3**). Examining treatment effects by means of a two sample t-test revealed that LT enhanced RPE associated neural responses in the left VS (peak Montreal Neurological Institute (MNI): x,y,z=-8,0,8, t_(59)_=4.24,k=243, P_FWE-cluster_<0.05, **Fig. 4a**) and bilateral orbitofrontal cortex (left OFC, peak MNI: x,y,z =-40,54,-16, t_(59)_=4.35, k=655, P_FWE-cluster_<0.05; right OFC, peak MNI, x,y,z=44,40,-14, t_(59)_=4.24,k=326, P_FWE-cluster_<0.05, **Fig. 4a**). Examination of extracted parameter estimates (spherical masks, radius: 6mm) revealed that these regions signaled positive but not negative RPEs under PLC whereas the LT-induced increase further enhanced positive RPE and instated negative RPE signaling in these regions (**Fig. 4b**). To further explore whether treatment differentially affected the RPEs in the VS we employed an independent mask from the Brainnetome atlas [38]. The use of an independent mask alleviates a potential bias of post hoc statistics [44] and the VS was chosen due to its critical involvement in RPEs. Analyzing extracted estimates from the VS further confirmed that losartan increased activation in the VS for both positive and negative RPEs (details **Fig. S7**).

**Fig. 4.**
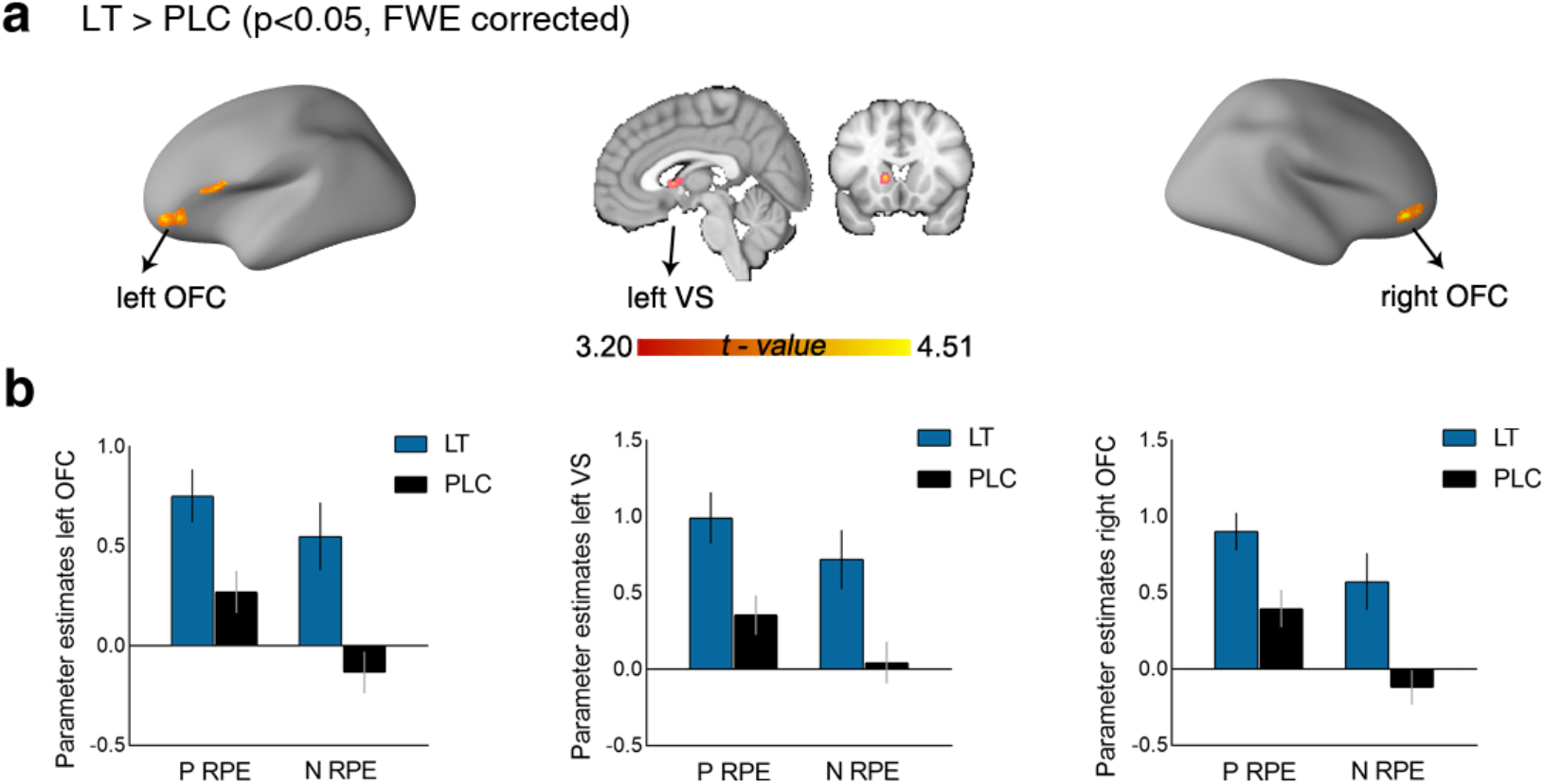
Losartan modulation on RPE-related neural response. **(a)** Comparison of losartan and placebo groups in RPE-related response suggested a losartan-triggered increased activation of bilateral orbitofrontal cortex and left ventral striatum. **(b)** For illustration purpose, parameter estimates extraction from spherical masks (radius: 6 mm) of identified left or right orbitofrontal cortex as well as left ventral striatum showed that losartan enhanced activation in these regions to RPE for both positive and negative outcomes. The error bars denoted standard error of the mean. LT, losartan; PLC, placebo. FWE-family-wise error, OFC-orbitofrontal cortex, VS-ventral striatum, PRPE/N-positive reward prediction error, NRPE-negative reward prediction error.

### LT sharpens differential neural representations for positive vs negative outcomes in the VS during early learning

We initially established an accurate whole brain multivariate predictive pattern for classifying positive and negative outcomes (accuracy, 89.34%, sensitivity and specificity, 88.52% and 90.16%, respectively **Fig. 5b**). Applying thresholding (bootstrapped 10,000 samples) and multiple comparisons correction (false discovery rate [FDR] corrected, p<0.001) revealed that a network including the VS, ventromedial prefrontal cortex, dorsomedial prefrontal cortex and middle frontal gyrus strongly contributed to the prediction of positive or negative outcomes during early learning (**Fig. 5a**, for a validation in an independent dataset see **Fig. S8**).

**Fig. 5.**
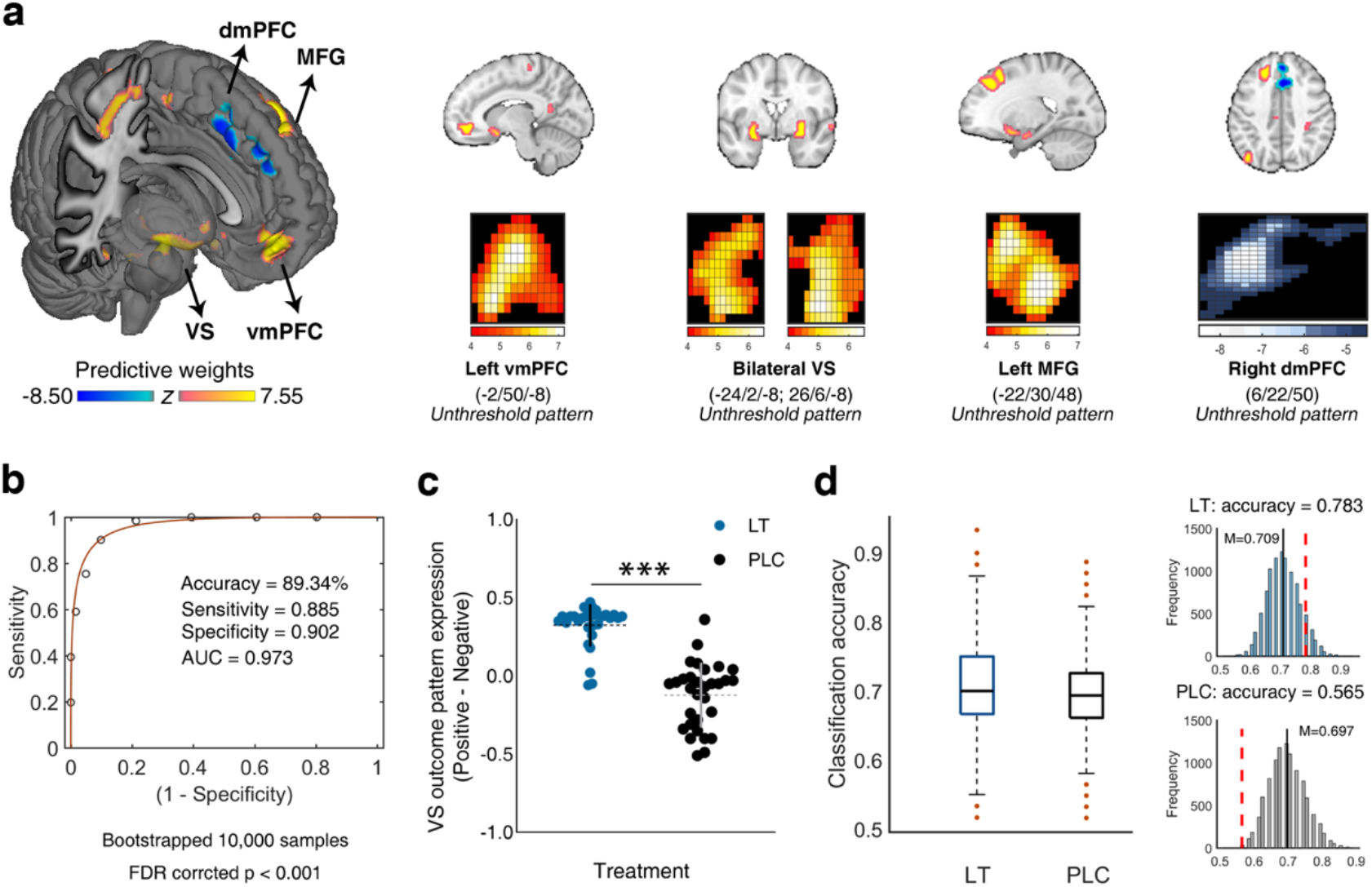
Multivariate neural predictive pattern results. **(a)** Neural predictive pattern consists of voxels in which activity reliably predicted positive versus negative outcomes during early phase of learning. This map shows weights that exceed a threshold (p<0.001, FDR corrected based on bootstrapped 10,000 samples) for display only. Hot color indicates positive weights and cold color indicates negative weights. **(b)** ROC plot. This neural predictive pattern yields a classification accuracy of 89.34% in a leave-one-subject-out cross validation procedure. **(c)** Losartan treatment increases the neural pattern of the ventral striatum for positive outcome. **(d)** In the losartan group a better classification performance relative to the placebo group was observed across all permutations. The histograms of classification accuracy for the ventral striatum neural expression for classifying positive and negative outcome from permutation tests are presented. The red line shows the true classification accuracy in the losartan and placebo group separately. The error bars denotes standard error of the mean and the black line shows the mean value. VS-ventral striatum; vmPFC-ventromedial prefrontal cortex; dmPFC-dorsomedial prefrontal cortex; MFG-middle frontal gyrus; AUC-area under curve; FDR-False discovery rate; LT-losartan; PLC-placebo; ***p<0.001.

Based on our a priori regional hypothesis about the crucial role of VS in reward learning we examined effects of LT on VS neural representations for positive outcomes. Our results suggested that only following LT - but not PLC - the VS expression accurately differentiated positive from negative outcomes (accuracy=78.33%, p<0.001, sensitivity=0.83, specificity=0.73, AUC=0.88; PLC, accuracy=56.45%, p=0.37, sensitivity=0.55, specificity=0.58, AUC=0.68), with a direct comparison between the treatment groups suggesting that LT specifically enhanced the VS representation for positive outcomes (t_(59)_=9.92,p<0.001,d=1.29, **Fig. 5c**). A group comparison on classification accuracy using permutation-inference further confirmed a significant treatment effect (t=-15.53, p<0.001, 95% CI, [-0.013, -0.010], **Fig. 5d**).

### LT accelerates choices and increased VS-dlPFC coupling for approaching maximum rewards during the transfer phase

In the transfer phase, both choosing A and avoiding B were significantly higher than chance level (50%) regardless of treatment (choose A: t=15.07, p<0.001, avoid B: t=15.16, p<0.001), which indicated most participants could successfully apply their previous learning experience in the transfer phase. The main effect of treatment on choice accuracy was not significant (β=-0.03, 95% HDI, [-0.13, 0.06]). However, analyzing choice reaction times revealed a significant main effect of choice behavior (β=-12.70, 95% HDI, [-23.18, -1.94]) and an interaction effect of treatment with choice behaviors (β=-27.36, 95% HDI, [-42.41, - 12.41]). Relative to PLC, LT accelerated choice times for choosing A (β=-83.52, 95% HDI, [- 147.03, -18.08], **Fig. 6a**), reflecting facilitated approach of the previously learned best option following LT. Furthermore, we explored how previous learning experience supported the optimal choice behaviors by conducting correlation analyses for learning rate and choice behaviors of transfer. We found that learning rate for negative outcomes was not significantly correlated with choice accuracy or response time of choose A or avoid B choices across treatment groups, yet in the LT group, subjects with high learning rate for positive outcomes during initial learning showed faster response towards maximum reward and avoiding worst stimulus in learning transfer (**Fig. 6b**).

**Fig. 5.**
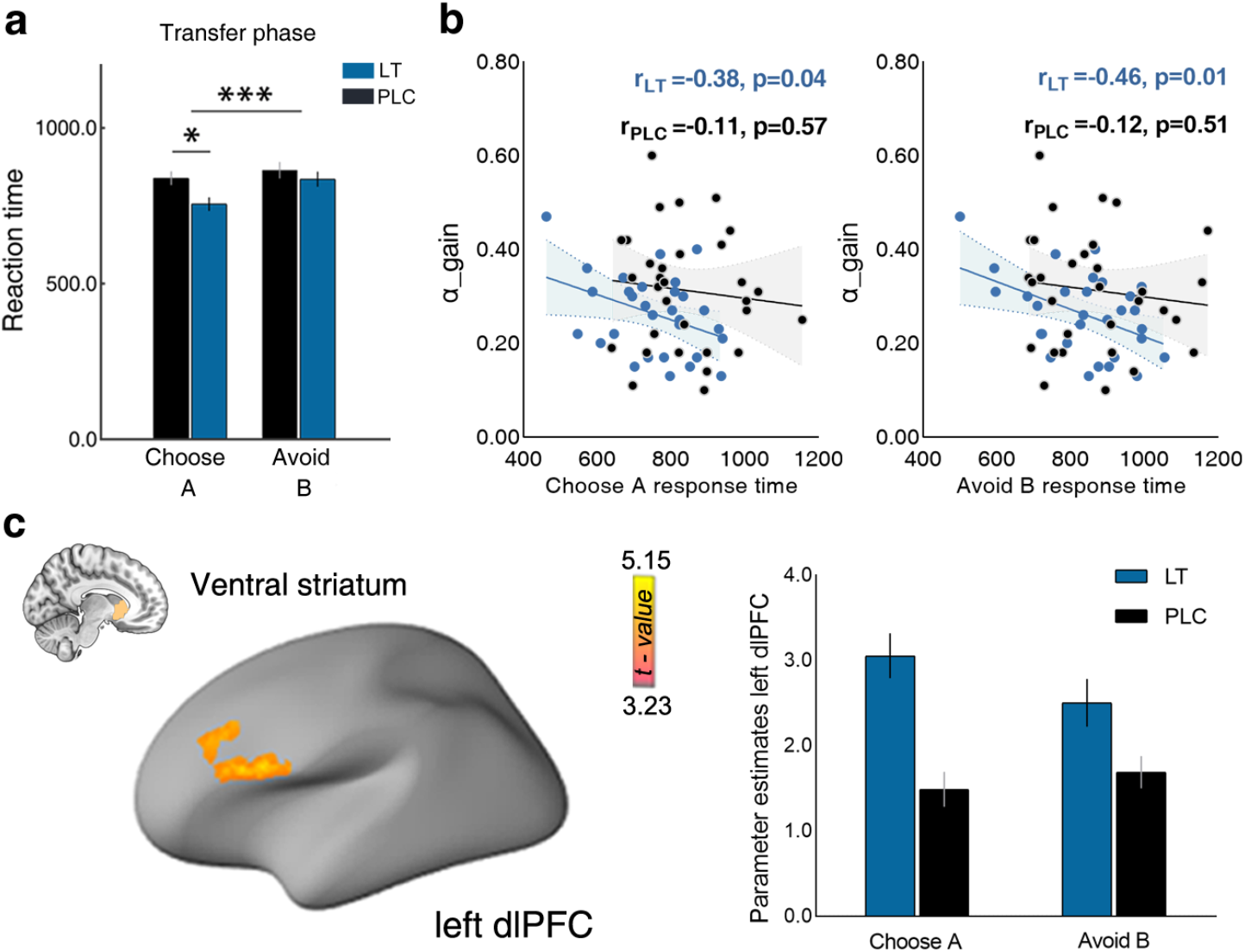
Behavioral and neural results for transfer phase. **(a)** In the transfer phase, all participants responded quickly when choosing stimulus A or avoiding stimulus B, and relative to the placebo group, losartan group exhibited faster responses for approaching stimulus A in a novel environment. **(b)** Moreover, for losartan-treated participants, higher learning rate for positive outcomes in initial learning phase was related to the accelerated responses toward maximum reward or avoiding worst option in transfer phase, while this association was abolished in placebo group. **(c)** For illustration purpose parameter estimates were extracted from a spherical (radius: 6 mm) ROI in the identified left dorsolateral prefrontal cortex (dlPFC) region. Losartan increased functional coupling between ventral striatum and left dlPFC when participants choose A stimuli in transfer phase. The statistical map of the left dlPFC was thresholded at p<0.001 uncorrected (whole-brain level) for display purpose. The error bars denoted standard error of the mean. LT-losartan; PLC-placebo.

On the neural activation level, we did not observe treatment effects of LT on learning transfer related regional activation, as well as on VS activity for each value difference level (all ps>0.05, **Fig. S9**). However, on the level of functional connectivity LT increased functional connectivity between the VS and left dorsolateral prefrontal cortex (left dlPFC, peak MNI: x,y,z=-48,22,28, t_(59)_=5.15, k=197, P_svc-FWEpeak_=0.01, **Fig. 6c**), reflecting an LT-induced enhancement of VS-dlPFC communication when approaching maximum rewards during learning transfer.

## DISCUSSION

The present pharmacological study utilized computational modeling in combination with fMRI to examine the effects of transient LT-induced AT1R blockade on reinforcement learning and the underlying neural mechanism in healthy individuals. On the behavioral level LT facilitated choice accuracy in the most difficult reward condition while it specifically reduced learning from negative outcomes and enhanced exploitatory choice behaviors. On the neural level, the behavioral effects were paralleled by regional-specific effects on ventral striatal-orbitofrontal reward systems, such that LT increased RPE signaling in these regions and sharpened the fine-grained neurofunctional distinction between positive and negative outcomes in the VS. During learning transfer, LT facilitated approach of the maximum rewarding option and enhanced VS-dlPFC functional connectivity. Overall, these findings indicate that LT-attenuated learning from negative feedback in the context of general positive outcome learning and a subsequent increased motivation to obtain maximum rewards during learning transfer, which on the neural level was accompanied by enhanced RPE and functional communication in fronto-striatal circuits.

We found that LT specifically enhanced choice accuracy for the most difficult condition suggesting that LT specifically improved learning under a low reinforcement probability. Computational modeling additionally allowed a more fine-grained examination of the behavioral effects by fitting trial-by-trial learning behavior and revealed that LT specifically reduced the learning rate for negative outcomes and enhanced exploitatory choices. The optimal learning ability could be understood when learning rate and other free parameters are considered simultaneously (e.g., explore-exploit tendency) in the RL model and the reward schedule [45]. Effects of LT on learning rate were outcome dependent, such that LT specifically decreased learning from negative outcomes, indicating an attenuated influence of negative information on reinforcement learning. Within the context of a stable reinforcement schedule, it is adaptive for an agent to ignore relatively rare and potentially misleading negative feedback given that an oversensitivity to negative outcomes would cause suboptimal choice behaviors. Therefore, decreased negative learning rate may signal the relatively high approach for positive outcomes in a stable reward contingency, which in turn may facilitate an exploitatory choice tendency in terms of consistent decisions for options with a higher expected reward [33]. Previous studies demonstrated that enhancing central dopaminergic activity increases choices towards monetary gains [1, 46]. The current pattern of results may reflect modulatory effects of RAS blockade on dopaminergic neurotransmission given that LT has been shown to induce stronger D1 receptor expression [47] which has been associated with better reward-associative learning [48]. These findings resonate with recent studies reporting an LT-induced enhancement of learning from positive relative to negative events [12] as well as an LT-induced shift from preferential social punishment towards social reward processing [17]. Together, this pattern of effects suggests that LT can attenuate the impact of negative information thus promoting motivation to select rewarding options.

On the neural level LT increased orbitofronto-striatal RPE-signaling and induced a more distinct neural expression for positive outcomes in the VS. The VS dopamine neurons are critically involved in RPE signaling and reward seeking [49], while the OFC is strongly implicated in computation of expected reward values and RPEs [50]. An LT-induced enhancement of the neural RPE signal and the representation of rewarding outcomes may reflect the potential for the RAS to modulate central dopaminergic neurotransmission during reinforcement learning. The AT1R is expressed densely in dopamine-rich brain areas [20], particularly in the striatum [51] and plays a key role in dopaminergic function [52]. Administering an antagonist of AT1R could mediate D1 density in the striatum [47] and block the functional response of the D2 receptor [21] - both of these receptors exhibit dense expression in ventral striatal and prefrontal regions crucially involved in reward learning [53, 54]. This may indicate a potential downstream effect of LT-induced AT1R blockade on DA signaling, in turn modulating reward learning within orbitofronto-striatal circuits, thus enhancing RPE encoding and reward representation in these regions.

During subsequent learning transfer, LT facilitated approach of the maximum reward in terms of accelerated decisions. This effect was not linked with the role of VS in value difference processing but was observed in the context of enhanced functional coupling of the VS with left dlPFC. Faster decisions for choosing the best options following LT may reflect an increased motivation to focus on maximizing rewards after reinforcement learning. The findings partly align with early research on dopaminergic modulation of reinforcement learning, which reported improved motivation for the highest-rewarding option during transfer, an effect that was explained as dopamine-dependent enhancement of learning signals [55]. The important role of fronto-striatal connectivity in reinforcement learning has been extensively documented [41], indicating that reward associations initially formed in the striatum are subsequently used to guide learning and decisions engaging the dlPFC [56]. To be specific, the dlPFC plays an important role in integrating and transmitting reward representations to the mesolimbic and mesocortical dopamine systems to initiate reward-motivated behaviors [57]. Reduced striato-dlPFC connectivity has been observed in in disorders characterized by a dysfunctional DA system [58] and linked with impaired reinforcement learning [59]. As such, the present findings of an LT-induced increase in VS-dlPFC connectivity when approaching rewards might reflect a modulatory role of angiotensin signaling on fronto-striatal communication via effects on dopaminergic circuits.

Given the repeatedly observed hypersensitivity for negative information and an increased impact of negative information on learning in depression [60], the current pattern of behavioral effects may reflect potential for LT to normalize biased processing in depression and in turn improve motivational deficits. The therapeutic potential in depression is further supported by early animal models suggesting a crucial role of the RAS in depression [61, 62] and documenting potential antidepressant behavioral effects of LT [63, 64]. Initial studies aimed at targeting reward processing and reinforcement learning impairments in depression via directly targeting the dopaminergic system [65, 66]. These studies revealed initially promising evidence for a therapeutic potential of DA agonist in depression including normalized neural functioning in fronto-striatal reward systems [65, 66] and anhedonia improvement [5]. However, effects on impaired reward learning were not observed and the clinical utility of DA agonist is limited by adverse effects such as triggering impulsive behaviors [67] and abuse [68]. The current pattern of results may point to a possibility that LT may represent a safe and potentially behavioral relevant strategy to modulate deficient reward learning and fronto-striatal functioning in depression.

While the current study found some evidence for a novel pathway to modulate reward learning, future studies are required to: (1) determine the potential of LT to influence reward learning and associated fronto-striatal deficits in depression, and (2) uncover the detailed interaction mechanism between the RAS with DA systems during reward learning such as incorporating receptor maps in combination with molecular imaging. In addition, future studies are required to demonstrate whether the observed effects generalize to women.

Taken together, we demonstrated that AT1R blockade via LT decreased negative learning rate but did not affect learning from positive outcomes, while increasing RPE signaling in orbitofronto-striatal regions and improving neural expression of positive outcomes in the VS. During the subsequent transfer, LT accelerated choices for maximizing rewards and increased VS-dlPFC functional coupling. Together, this pattern may reflect a promising mechanism of LT as a potential treatment to normalize impaired reward learning and fronto-striatal functioning in depression.

## ACKNOWLEDGMENTS

We truly thank all colleagues for the impressive discussions that help us to improve the manuscript and the help during the FMRI data acquisition, and all volunteers that participated in our study. This work was supported by the National Natural Science Foundation of China (Grant No. 32250610208), China MOST2030 Brain Project (Grant No. 2022ZD0208500) and the National Key Research and Development Program of China (Grant No. 2018YFA0701400).

## AUTHOR CONTRIBUTIONS

TX and BB designed the study. TX, LW, GJ, RZ conducted the experiment and collected the data. TX, LJ, XZ, FZ performed the data analysis. TX and BB wrote the manuscript draft. JK, CZ, FZ, WZ, YS and BB critically revised the manuscript draft.

## COMPETING INTERESTS

The authors report no biomedical financial interests or potential conflicts of interest.

## DATA AND MATERIALS AVAILABILITY

Unthresholded group-level statistical maps are available on NeuroVault (https://neurovault.org/collections/12001/). Additional data related to study is available from the corresponding author upon reasonable request. A preprint of the manuscript was archived on the biorxiv.org repository (doi: https://doi.org/10.1101/2022.03.14.484364).The present study was pre-registered on Clinical Trials.gov (Trial name: The effects of losartan on reward reinforcement learning; Registration number: NCT04604938; URL: https://clinicaltrials.gov/ct2/show/NCT04604938).

## Notes

### Competing Interest Statement

The authors have declared no competing interest.

### Summary of Updates

Revision including further analyses and extended discussions

