## Supplementary material for "Angiotensin blockade enhances motivational reward learning via enhancing striatal prediction error signaling and frontostriatal communication": supplental

**Affiliations**
[1] The Center of Psychosomatic Medicine, Sichuan Provincial Center for Mental Health, Sichuan Provincial People's Hospital, University of Electronic Science and Technology of China, Chengdu, China
[2] MOE Key Laboratory for Neuroinformation, School of Life Science and Technology, University of Electronic Science and Technology of China, Chengdu, China
[3] Department of Psychology, University of Cambridge, Cambridge, UK
[4] Behavioural and Clinical Neuroscience Institute, University of Cambridge, Cambridge, UK
[5] Max Planck School of Cognition, Leipzig, Germany
[6] Faculty of Psychology, Southwest University, Chongqing, China
[7] Key Laboratory of Cognition and Personality, Ministry of Education, China

***Correspondence to**
Benjamin Becker: https://orcid.org/0000-0002-9014-9671
University of Electronic Science and Technology
Xiyuan Avenue 2006, 611731 Chengdu, China
mail:

### Supplemental Methods

#### Participants

Seventy right-handed healthy male participants from a local university were enrolled and screened according to following criteria: 1) no current or a history of psychiatric, neurological, or other medical disorders, 2) no current or regular use of psychotropic substances including nicotine, 3) a body mass index<18 or >24.9, 4) no visual or motor impairments, and 5) no contraindications for MRI or losartan, 6) refrain from excessive caffeine before the study.

#### Potential confounders

To control potential group differences in pre-treatment mood and anxiety as well as treatment effects on current mood and anxiety the Positive and Negative Affect Schedule and the Spielberger State-Trait Anxiety Inventory were administered before treatment and after the experiment. The Beck Depression Inventory was additionally administered before treatment. In addition, potential treatment effects on working memory capacity and attention were assessed by means of a 2-back working memory task and a digit cancellation test. Although previous studies suggested that the effects of LT on cardiovascular activity only manifest after at least 3 hours after p.o. administration while the central effects occurred faster (Goldberg et al., 1993; Mechaeil et al., 2011), the heart rate and blood pressure were monitored before and after the treatment administration, as well as after the fMRI experiment. Finally, the transfer phase employed new pairings of the stimuli, leading to a total of six value difference levels among all stimulus pairs (e.g., C vs E (70% vs 60%) or C vs D (70% vs 30%)). The comparably large rewarding probability or value difference of the re-organized stimulus pairs may lead to a bias of treatment effects on choice accuracy and response time. To further explore a potential influence, we conducted two mixed ANOVA models with treatment and value difference level as independent variables, and choice accuracy or response time as dependent variable, respectively. The analyses did not yield significant main or interaction effects involving treatment (**Fig. S2, Table S1-S2**) arguing against a confounding effect of value difference levels.

#### Multilevel Bayesian linear model analyses

We employed a multilevel Bayesian linear model to analyze trial-by-trial behavior via using a Bayesian regression model in the Stan (brms) package (Bürkner, 2017) in R. For the learning behavior, the fixed and random trial-by-trial effects were coded with a varying intercept on per subject basis. Specifically, the choice accuracy for choosing the better stimuli in a pair was the dependent variable (correct =1, wrong =0) whereas the stimulus pair was set as the within-subjects variable ranging from easiest to the most difficult (AB=1, CD=0, EF=-1). The binary covariates of interest treatment (LT=1, PLC=0) and fMRI runs (Run1=1, Run2=2) were also included as between-subjects variables.

For the test phase, consistent with previous reinforcement learning studies (Erin C. Dowd et al., 2016; Jocham et al., 2011; Simon et al., 2010), we also explored the differential performance in selecting the best and avoiding the worst stimulus within the novel pairings. As such the regression models during the transfer phase focused on trials during which either the A or B stimulus were presented. The within-subject variable was therefore changed to the approach A or avoid B variable (A=1, B=-1) on each trial while the dependent variable was the choice accuracy in choosing A in approach A trials or not choosing B in avoid B trials (correct=1, incorrect=0). Treatment (LT=1, PLC=0) was also included as between-subject variable with a varying intercept per subject.

For these regression models samples of each parameter's posterior distribution were drawn with a Hamiltonian Monte Carlo sampling algorithm implemented in Stan (Carpenter et al., 2017; Hoffman & Gelman, 2014). The model was fitted using three independent Markov Chains, each with 5000 iterations of which 2000 were warm-up to calibrate the sampler, leading to a total of 9000 posterior samples. To confirm that the samples for each chain converged to the same posterior distribution, the Geweke diagnostic and R-Hat statistics were calculated (Gelman & Rubin, 1992). Finally, we reported main effects of treatment, stimulus pair, fMRI runs, and interaction effects of stimulus pair with treatment on credibly different parameters, i.e. when more than 95% of the posterior distribution is above/below zero.

**Computational modeling**

We explored the learning rate from participants' choice behavior by using a computational reinforcement learning 'Q-learning' algorithm with a hierarchical Bayesian parameter estimation for data collected in the separate fMRI runs. In hierarchical models, group and individual parameter distributions are fitted simultaneously and constrain each other, leading to greater statistical power over standard non-hierarchical methods (Pedersen & Frank, 2020). In the current study, the model estimates the expected value (Q) of stimulus (A-F) based on the individual sequences of choices and experienced outcomes. The expected value for choosing one stimulus in a pair will be updated according to the following equation:

$$Q_{cs}(t+1) = Q_{cs}(t) + \begin{cases} \alpha_{Gain}[r(t) - Q_{cs}(t)], & \text{if} \quad r = 1 \\ \alpha_{loss}[r(t) - Q_{cs}(t)], & \text{if} \quad r = 0 \end{cases}$$

Where the term $Q_{cs}(t+1)$ is the expected value of chosen stimulus on trial t + 1, which is determined by the expected value of last chosen stimulus on trial t and the learning rate. Because previous LT work demonstrated a distinguishable learning trend for positive and negative outcomes (Pulcu et al., 2019), we modeled separate learning rate parameters for positive (gain) and negative (loss) reward prediction errors (RPE, $r(t)-Q_{cs}(t)$). The choice on trial t followed by a positive feedback would be weighted by the learning rate ($0 \leq \alpha_{Gain} \leq 1$) for positive reward prediction error (PRPE), or it was weighted by the learning rate ($0 \leq \alpha_{Loss} \leq 1$) for negative reward prediction error (NRPE) if followed by a negative feedback. The learning rate scales the degree to which the prediction error is used to update expected values, and a higher learning rate reflects a faster value updating relying on recent rewards whereas a lower learning rate indicates gradual value update uncovering long-lasting effects of outcome. To avoid the initial bias in the expected value the Q value were all set to the 0.5 before learning. In each stimulus pair, the probability of choosing the object stimuli over the other one is estimated using the softmax rule as follows:

$$P_{cs}(t) = \frac{\exp(\beta \times Q_{cs}(t))}{\exp(\beta \times Q_{ucs}(t)) + \exp(\beta \times Q_{cs}(t))}$$

The parameter β with a range between 1 to 15 is referred to as inverse temperature revealing the exploratory or exploitatory choice behavior. High temperature β indicates a consistent choice behavior with the stimulus of higher expected value being invariably selected, whereas low estimates of this parameter reflects random choice behavior irrespective of the expected value of the stimulus. The parameters $\alpha_{gain}$, $\alpha_{Loss}$ and temperature β were estimated for each subject and each run, being modified to maximize the probability of the actual choices under the model.

This model was estimated by using Markov Chain Monte Carlo (MCMC) inference. Models were implemented using the Stan programming language (Carpenter et al., 2017; Hoffman & Gelman, 2014). We ran three chains of 5000 samples each (discarding the first 2500 of each chain for burn-in), and ensured convergence using manual examination of the trace plots (hairy caterpillars, easily moving around the parameter space) and evaluation of R-Hat statistics, which were all less than 1.1 (Gelman & Rubin, 1992). In addition group-level parameter estimate distributions were obtained (**Fig. S1a**) and an "optimism bias" that revealed a higher learning rate for positive versus negative outcomes was observed, which is in line previous studies using separate learning for positive and negative events (Jahfari et al., 2019; Jahfari et al., 2020; Lefebvre et al., 2017). To check whether the model sufficiently captured actual choice behavior of participants, we simulated the probability of choosing the best option using the posterior distributions of the fitted free parameters of each participant. The results revealed that the model showed good representation of overall learning across each group and stimulus pair (**Fig. S1b**). We finally estimated the Final Q values across each stimulus and group from the fitted model, finding the model captured the declining Q value according to decreasing reward contingency well (**Fig. S1c-1d**).

We additionally considered another model with only one learning rate for updating expected values of both positive and negative outcomes. The model comparison was conducted by using the Bayesian information criterion (smaller value indicates a better fit) (Schwarz, 1978) on an individual subject level and the Bayesian model comparison on the population level to prove the better representation of the chosen model to the original data. In addition, we also compared these two models by means of the leave-one-out cross-validation information criterion (LOOIC) procedure. LOOIC is highly recommended for model comparisons of hierarchical Bayesian structures that use MCMC sampling (Vehtari et al., 2017). LOOIC estimates point-wise out-of-sample prediction accuracy from a fitted Bayesian model using the log-likelihood evaluated at the posterior simulations of the parameter values and could be extracted using the "loo"package in R (Ahn et al., 2017; Vehtari et al., 2017; Yao et al., 2018). We found that the model with a single learning rate did not explain choice behavior better than the model with dual learning rates (Mean $BIC_{(\alpha, \beta)}$=108.58, Mean $BIC_{(\alpha Gain, \alpha Loss, \beta)}$=102.51, $t_{(60)}$=4.92, p<0.001,95%CI,[3.60,8.54]; $Model_{(\alpha, \beta)}$ BIC=120.91, $Model_{(\alpha Gain, \alpha Loss, \beta)}$ BIC=114.84; $Model_{(\alpha, \beta)}$ LOOIC±SE= 5914.30±247.20, $Model_{(\alpha Gain, \alpha Loss, \beta)}$ LOOIC±SE=5580.81±258.80). Therefore, the model with three free parameters $\alpha_{Gain}$, $\alpha_{Loss}$, and β (as presented in the manuscript) was determined as the best fitting model.

**Mediation analyses**

In order to test the relationship between learning parameter such as expected value, treatment and choice accuracy in learning phase, we performed exploratory mediation analyses using the Mediation Toolbox (https://github.com/canlab/MediationToolbox). The mediation analysis tests whether the observed covariance between an independent variable (X) and a dependent variable (Y) could be explained by a third variable (M). Significant mediation effect is obtained when inclusion of M in a path model of the effect of X on Y significantly alters the slope of the X ‑ Y relationship. That is, the difference between total (path c) and direct (non-mediated, path c′) effects of X on Y (i.e., c−c′), which could be performed by testing the significance of the product of the path coefficients of path a × b,is statistically significant. Here we examined whether the expected value in each stimulus pair during learning phase mediated the effects of treatment on choice accuracy in corresponding stimulus pair. We used bias-corrected accelerated bootstrapping (10,000 replacements) for significance testing. However we only found that expected value sensitivity played a mediating role in the effects of treatment on choice accuracy in the hardest stimulus pair (**Fig. S5**).

**MRI data acquisition and preprocessing**

MRI data were collected on a 3.0 Tesla system (GE MR750, General Electric Medical System, Milwaukee, WI, USA). High-resolution brain anatomical MRI images were acquired using a T1-weighted sequence (TR = 6ms; TE = 2ms; flip angle = 9°; field of view = 256 × 256 mm; matrix size = 256 × 256; voxel size = 1 × 1 × 1 mm; number of slices, 156; slice thickness = 1 mm) to improve spatial normalization of the functional data and exclude subjects with apparent brain pathologies. Functional data using blood oxygenation level-dependent (BOLD) contrast were obtained using a T2*-weighted echo planar imaging sequence (TR = 2000ms; TE = 30ms; slice number, 39; slice thickness = 3 mm; flip angle = 90°; field of view = 240 × 240 mm; voxel size = 3.75 × 3.75 × 4 mm; resolution = 64 × 64).

All fMRI images were preprocessed and analyzed using standard procedures in SPM 12 (Statistical Parametric Mapping; http://www.fil.ion.ucl.ac.uk/spm/; Wellcome Trust Centre for Neuroimaging). The first 5 volumes of each functional time series were discarded to allow for T1 equilibration. Remaining images were corrected for acquisition time delay, realigned to correct for head motion, unwarped for magnetic field inhomogeneities correction, and co-registered with the T1-weighted structural image. After that the images were normalized to Montreal Neurological Institute (MNI) standard space (interpolated to 2 × 2 × 2 mm voxel size) using the segmentation parameters from the anatomical images, and were then spatially smoothed using an isotropic Gaussian kernel with full-width at halfmaximum (FWHM) of 8 mm.

**Multi-Voxel Pattern analysis (MVPA)**

We used a linear support vector machine (C=1) implemented in Canlab core tools (https://github.com/canlab/CanlabCore) to develop a multivariate pattern classifier for positive and negative outcome. The pattern classifier was trained on subject-wise univariate positive outcome>baseline and negative outcome>baseline contrasts during the early learning phase. To avoid overfitting, the classification performance was evaluated by a leave-one-out cross validation procedure (LOOCV). The resulting cross-validated neural pattern of outcome consists of the weights of each voxel in predicting the positive or negative outcome presentation plus the intercept. The pattern expression values were generated by taking the dot-product between the whole-brain unthresholded classifier weights with participant-specific brain activity maps. Moreover, the pattern expression estimates for each individual were obtained from patterns trained on other participants' data to avoid circularity. Given that theory and experiments also outline the LOOCV may lead to unstable and biased estimates (Varoquaux et al., 2017), we also validate our results by conducting the same analysis in another independent placebo data that was collected by using a relatively similar probabilistic reinforcement learning task in our group. Specifically, we first established an accurate whole brain multivariate predictive pattern for classifying positive and negative outcomes (**Fig. S6**) and obtained a highly similar neural pattern in the independent dataset of subjects treated with p.o. placebo. Subsequently we applied the neural pattern classifier from our study to the independent placebo dataset and further demonstrated excellent performance of this classifier (accuracy, 90.41%, sensitivity and specificity for different outcomes, 81.58% and 84.21%).

Based on our a priori regional hypothesis and the key role of the ventral striatum in the reward processing, the analysis of LT effects on the expression focused on the ventral striatum. To this end we identified effects of LT on the ventral striatum outcome expression by examining the treatment difference on the classification accuracy of ventral striatum pattern in differentiating positive versus negative outcome with permutation-based inferences (10000 permutations). The ventral striatum included the ventral caudate and nucleus accumbens (Zhou et al., 2019).

**General psychophysiological interaction analysis**

To investigate the effects of losartan on the functional coupling between the ventral striatum and the frontal cortex during the transfer phase, we performed a general psychophysiological interaction (gPPI) analysis. The gPPI approach allows to accommodate more than two experimental conditions while facilitating a greater sensitivity and specificity compared to conventional PPI implementations in SPM (Mclaren et al., 2012). On the first level the model included separate conditions corresponding to the events of approaching A and avoiding B choices and the respective PPI regressor terms as well as one regressor modelling all other choices (e.g., choose E in EF stimulus pair) and the six head motion parameters as covariates of no interest. Given that animal and human studies largely indicated the interaction of striatum with the prefrontal regions as a key mechanism for reinforcement learning (Averbeck & O'doherty, 2021; Lowet et al., 2020), we specifically modelled functional connectivity from the ventral striatum serving as seed region to all voxel within the prefrontal cortex (determined by the Brainnetome atlas (Fan et al., 2016)).

### Supplemental Results

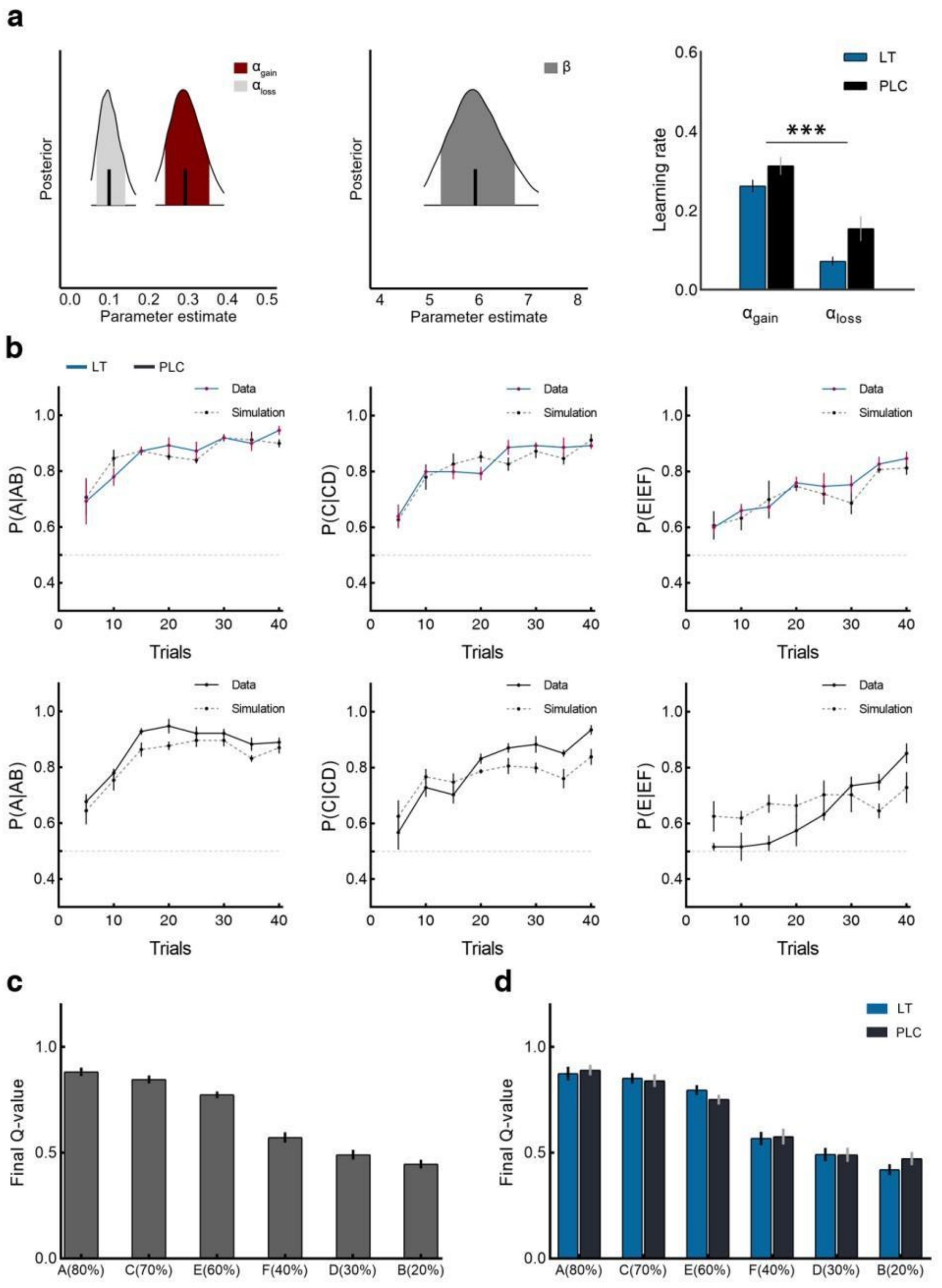


**Fig. S1 Computational model assessment and simulation. (a)** Group-level parameter estimate distributions. Similar to other studies using dual learning rates, higher learning rates for positive compared to negative outcome was observed, termed “optimism bias” (Jahfari et al., 2019; Jahfari et al., 2020; Lefebvre et al., 2017). This optimism bias was also observed for the individual parameter estimates, with higher learning rate for positive outcome in both groups. **(b)** Simulation of the fitted model. The probability of choosing the best option in each stimulus pair was simulated by using the posterior distributions of the fitted model parameters of each subject.

The plots of the model against observed data across each treatment group and stimulus pair revealed that the fitted model showed a good representation of the overall learning. **(c-d)** Final Q value across each stimulus in whole group, and in each treatment group, which showed that the fitted model did a good job in capturing declining Q value according to decreasing reward probability. The error bars denoted standard error to the mean. LT, losartan; PLC, placebo.

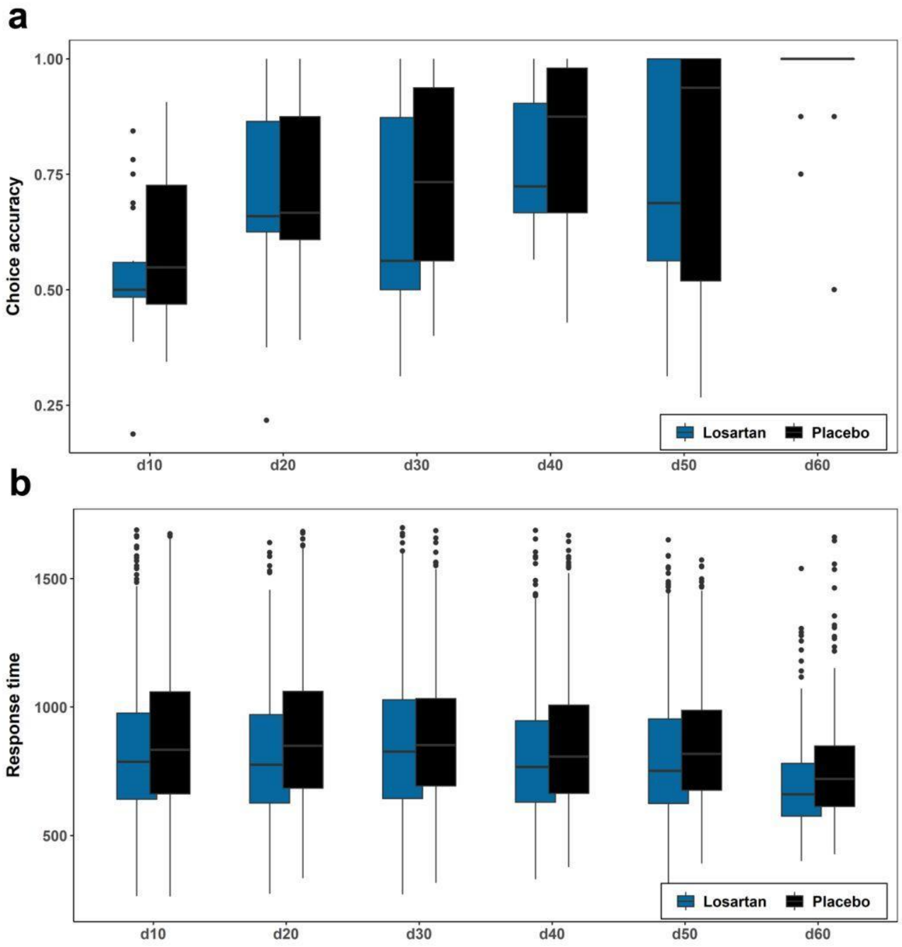


**Fig. S2 Effects of rewarding probability differences and treatment on choice behaviors in transfer phase. (a -b)** Losartan and placebo-treated subjects showed comparable choice accuracy and response time towards stimulus pairs with distinct rewarding probability. d10 - 60 indicates the six level of rewarding probability differences.

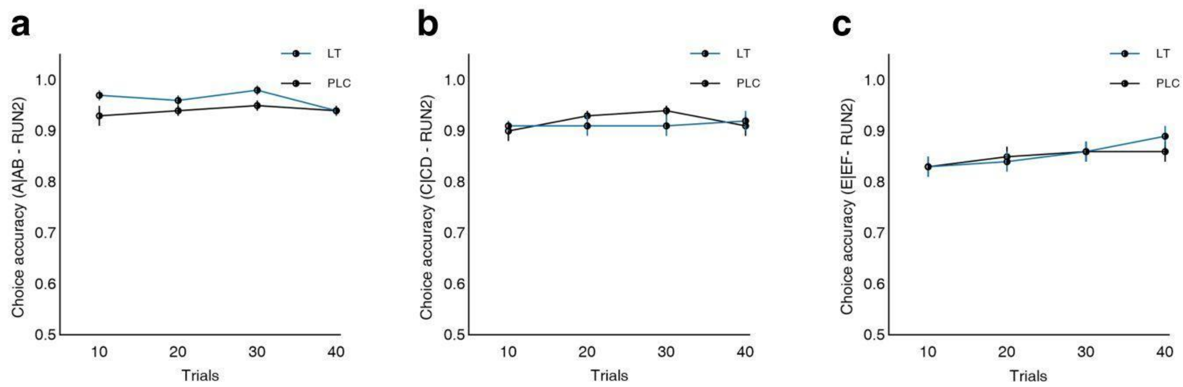


**Fig. S3 The raw learning curve for trial-by-trial choice accuracy across stimulus pairs in second fMRI run. (a-c)** Subjects showed constantly high choice accuracy during the learning for the second run, the latter was possibly driven by the practice leading to a ceiling effect effects. The error bars denoted standard error to the mean. LT, losartan; PLC, placebo.

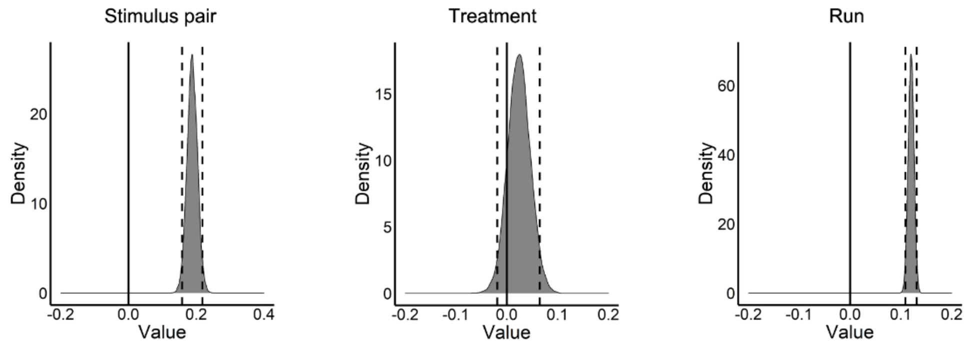


**Fig. S4 Bayesian regression model results.** We observed a significant main effect of stimulus pair such that participants successfully learned to choose better option from the easy stimulus pair. However no main effect of treatment was found, while the main effect of fMRI runs reached statistical significance such that choice accuracy in the second fMRI run showed almost ceiling effects thereby being higher than that in the first run.

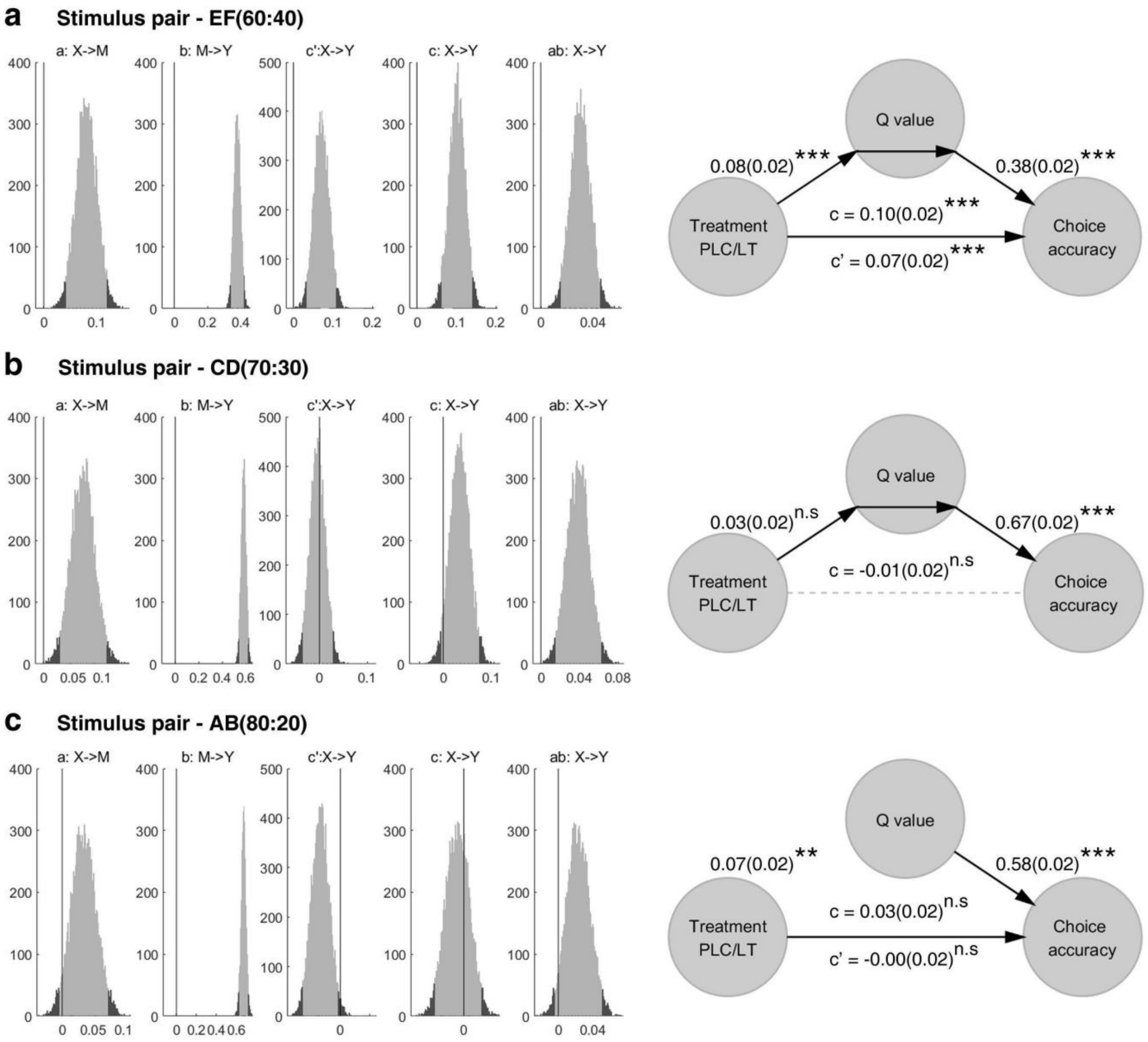


**Fig. S5 Mediation results for different stimulus pair. (a)** Mediation analysis results shows that expected value sensitivity (denoted as Q value) mediates the effects of losartan on choice accuracy for hardest stimulus pair. **(b-c)** The expected value could not mediate losartan effects on choice accuracy for easy or the easiest stimulus pair. n.s indicates non-siginficant, **p<0.01, ***p<0.001 (bootstrap tests; two-sided; uncorrected)**.**

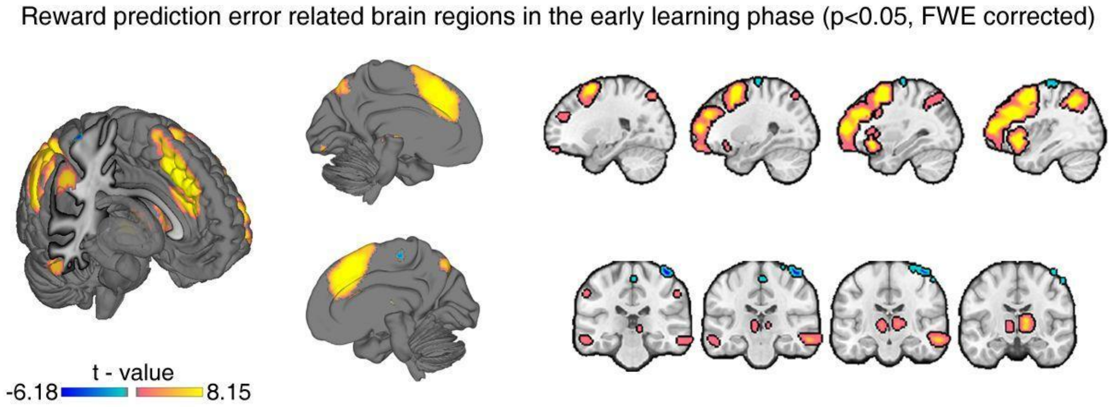


**Fig. S6 Reward prediction error related brain regions in the early learning phase.** Concerning the outcome onset for the early learning phase (first FMRI run), we found a neural network including the superior medial frontal gyrus, middle cingulate cortex, insula, thalamus, caudate, inferior parietal cortex, supramarginal area, middle temporal gyrus and inferior temporal gyrus, as well as the fusiform positively encoding reward prediction error (RPE), and the precentral and supplementary motor areas negatively encoding the RPE ($P_{FWE\text{-}peak}<0.05$).

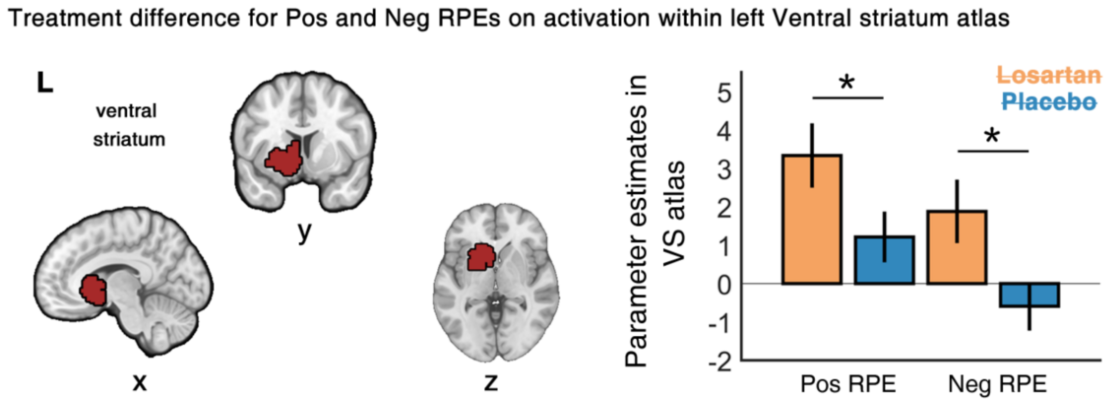


**Fig. S7 Treatment difference for positive (Pos) and negative (Neg) reward prediction errors (RPEs) on activation within independently defined atlas of ventral striatum (VS)**. We further explored differential treatment effects on the positive and negative RPEs using an atlas-based mask for the VS and extracted the corresponding parameter estimates for the Pos RPE and Neg RPE from each treatment group. Post hoc analyses revealed that losartan enhanced left VS activation related to both, Pos and Neg RPEs as compared to placebo (Pos RPE, $t_{(59)}=1.99$, Cohen's d=0.50; Neg RPE, $t_{(59)}=2.38$, Cohen's d=0.59, all ps<0.05).

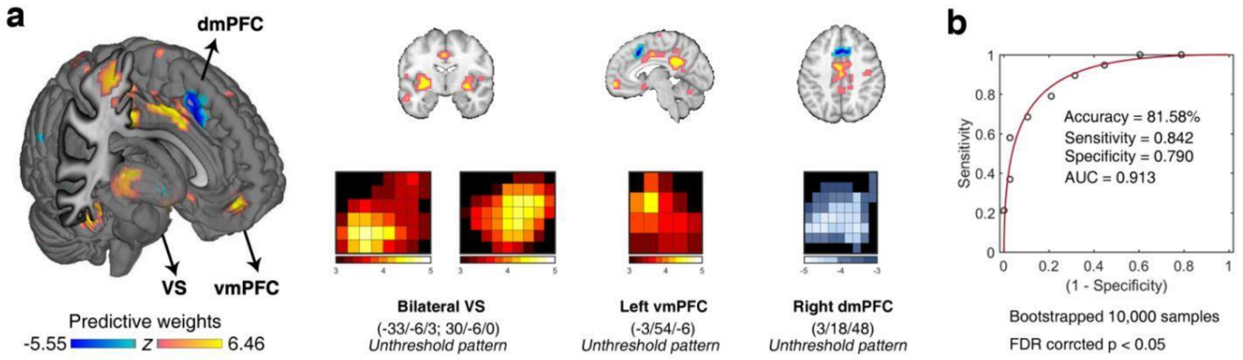


**Fig. S8 Multivariate neural predictive pattern results**. **(a)** Neural predictive pattern consists of voxels in which activity reliably predicted positive outcome versus negative outcome during learning in another probabilistic reinforcement learning task. The map shows weights that exceed a threshold (p<0.05, FDR corrected based on bootstrapped 10,000 samples) for display only. Hot color indicates positive weights and cold color indicates negative weights. **(b)** ROC plot. The neural outcome-predictive pattern yielded a classification accuracy of 81.58% in a leave-one-subject-out cross-validation procedure.

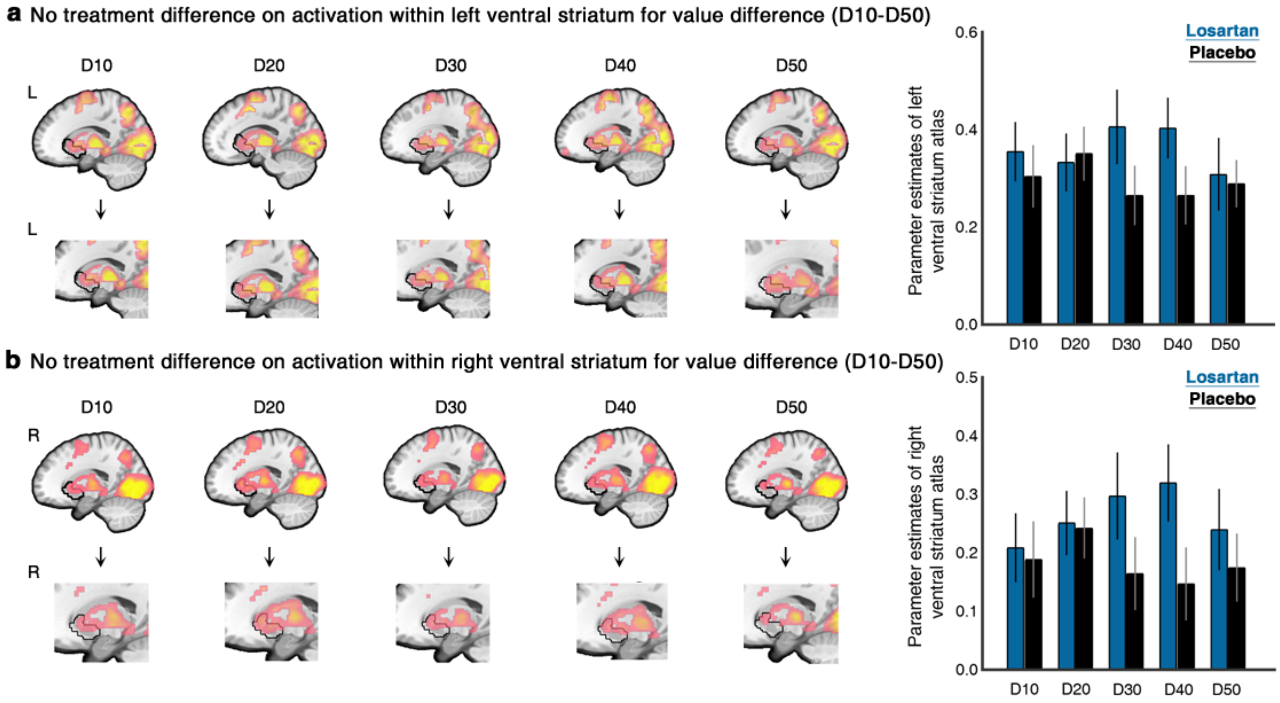


**Fig. S9 No treatment difference was observed on activation within the left (a) and right (b) ventral striatum (atlas mask) for each value difference level**. The D indicates the value difference of stimulus pairs in learning transfer phase.

**Table S1. Effects of rewarding probability differences and treatment on choice accuracy in transfer phase**

| Variables | β estimate | Bayesian 95% HDI |
|---|---|---|

| | | |
|---|---|---|
| Rewarding probability differences | 0.06 | [0.05, 0.07] |
| Treatment | -0.06 | [-0.14, 0.03] |
| Rewarding probability differences*Treatment | 0.00 | [-0.01, 0.02] |

**Table S2. Effects of rewarding probability differences and treatment on response time in transfer phase**

| Variables | β estimate | Bayesian 95% HDI |
|---|---|---|
| Rewarding probability differences | -16.29 | [-20.68, -12.16] |
| Treatment | -72.20 | [-146.62, 5.55] |
| Rewarding probability differences*Treatment | 4.97 | [-1.01, 11.01] |

**Table S3. Reward prediction error related brain regions in the first fMRI run ($p_{FWE\text{-}peak}$<0.05)**

| | | MNI coordinates | | | | | |
|---|---|---|---|---|---|---|---|
| **Region** | **side** | **x** | **y** | **z** | **$T_{(60)}$** | **k** | **$p_{FWE\text{-}peak}$** |
| *--Positively encoded RPE* | | | | | | | |
| Superior medial frontal gyrus | L/R | 0 | 32 | 40 | 10.87 | 19927 | <0.001 |
| Middle cingulate cortex | R | 6 | 28 | 32 | 10.71 | | <0.001 |
| Insula | L | -32 | 22 | -2 | 10.64 | | <0.001 |
| Thalamus | R | 12 | -8 | 8 | 7.83 | 500 | <0.001 |
| Thalamus | R | 8 | -12 | -2 | 7.25 | | <0.001 |
| Thalamus | L | -10 | -18 | 4 | 6.01 | | 0.002 |
| Thalamus | L | -14 | -6 | 6 | 5.02 | 1 | 0.040 |
| Caudate | L | -10 | 8 | 4 | 7.21 | 385 | <0.001 |
| Caudate | R | 12 | 4 | 8 | 7.71 | | <0.001 |
| Pallidum | L | -14 | 0 | -2 | 6.77 | | <0.001 |
| Pallidum | R | 16 | 4 | 4 | 5.69 | 19 | 0.005 |
| Pallidum | R | 18 | 2 | -4 | 5.08 | | 0.034 |
| Putamen | R | 32 | 14 | -4 | 4.99 | 1 | 0.045 |
| Inferior parietal cortex | L | -54 | -36 | 44 | 9.2 | 3705 | <0.001 |

| | | | | | | | |
|---|---|---|---|---|---|---|---|
| Inferior parietal cortex | L | -40 | -54 | 50 | 9.11 | | <0.001 |
| Inferior parietal cortex | R | 40 | -50 | 50 | 9.03 | | <0.001 |
| Superior parietal cortex | R | -28 | -62 | 46 | 7 | | <0.001 |
| Supramarginal | R | 50 | -44 | 32 | 9.06 | 3686 | <0.001 |
| Supramarginal | R | 58 | -50 | 26 | 8.41 | | <0.001 |
| Middle temporal gyrus | R | 58 | -20 | -14 | 7.15 | 404 | <0.001 |
| Middle temporal gyrus | L | -46 | -58 | 0 | 5.37 | | 0.014 |
| Middle temporal gyrus | L | -58 | -28 | -14 | 5.87 | 97 | 0.003 |
| Inferior temporal gyrus | R | 60 | -30 | -18 | 5.66 | | 0.005 |
| Inferior temporal gyrus | R | 58 | -38 | -28 | 5.13 | | 0.029 |
| Fusiform | L | -44 | -62 | -16 | 6.13 | 217 | 0.001 |
| Fusiform | R | 44 | -56 | -20 | 5.46 | 52 | 0.010 |
| Fusiform | R | 40 | -46 | -24 | 5.04 | 5 | 0.038 |
| Calcarine | L | -2 | -86 | -12 | 5.48 | 24 | 0.009 |
| --*Negatively encoded RPE* | | | | | | | |
| Precentral | R | 40 | -26 | 70 | 7 | 152 | <0.001 |
| Precentral | R | 42 | -18 | 68 | 6.18 | | 0.001 |
| Precentral | R | 26 | -20 | 74 | 5.27 | | 0.019 |
| Precentral | R | 54 | -14 | 58 | 5.36 | 12 | 0.014 |
| Supplementary motor area | R | 2 | -24 | 60 | 5.35 | 19 | 0.015 |

MNI, Montreal Neurological Institute; L, left; R, right.
